# Evolutionary potential of the *Escherichia coli* antimutator Δ*nudJ* is reduced via altered mutational spectrum

**DOI:** 10.1101/2025.01.28.635215

**Authors:** R Green, H Richards, D Ozbilek, F Tyrrell, V Barton, SC Lovell, DR Gifford, M Lagator, AJ McBain, R Krašovec, CG Knight

## Abstract

The rate of spontaneous mutation is a key factor in determining the capacity of a population to adapt to a novel environment, for example a bacterial population exposed to antibiotics. Genetic and environmental factors controlling the mutation rate commonly also cause shifts in the relative rates of different mutational classes, i.e. the spectrum of spontaneous mutations. When the mutational spectrum is altered, the relatively enriched and depleted mutations may differ in their fitness effects. Here we aim to explore how a reduced mutation rate and altered mutational spectrum can contribute to adaptation in *Escherichia coli*. We measure mutation rates across a set of Nudix hydrolase deletants, finding multiple strains with an antimutator phenotype. We focus on the antimutator Δ*nudJ* which can cause a 6-fold mutation rate reduction relative to the wildtype with an altered mutational spectrum. Not only does *nudJ* deletion reduce the probability of antibiotic resistance arising but, in the case of rifampicin, the accessible resistance mutations under this altered mutation spectrum have greater fitness costs. While this effect is, so far, specific to this drug, it opens up the possibility of reducing the likelihood of resistance mutations establishing through simultaneous modification of mutation rate and spectrum. The identification and characterisation of antimutator alleles such as Δ*nudJ* provides potential targets for ‘antievolution drugs’ which could be applied during antibiotic treatment to inhibit NudJ activity. Such an approach could suppress spontaneous resistance evolution both through fewer resistance mutations and by limiting access to the fittest mutations.

## Introduction

Understanding what controls microbial mutation rates is a central question in microbial evolution with important applications for tackling antimicrobial resistance and improving the reliability of engineered microbes. Rapid adaptation by ‘mutator’ strains, with elevated mutation rates, can allow bacterial populations to overcome single (Couce et al. 2015), sequential (Elgrail et al. 2024) and double (Gifford et al. 2023) antibiotic therapy. Conversely ‘antimutator’ strains, with reduced mutation rates, are less prone to developing antimicrobial resistance (Ragheb et al. 2019) and can provide more sTable Sengineered strains for synthetic biology applications (e.g. (Pośfai et al. 2006; Umenhoffer et al. 2010; Csörgő et al. 2012; Deatherage et al. 2018; ScarabGenomics 2024)). Identifying ways to reduce bacterial mutation rates is therefore an area of great interest.

Increased mutation rates, caused by the disruption of genes involved in DNA replication and repair, are observed relatively frequently in both laboratory grown bacteria (e.g. (Good et al. 2017; Callens et al. 2023)) and clinical infections (e.g. (Oliver et al. 2000; Couce et al. 2016)). In contrast, strains with genetic reduction of mutation rates, termed ‘antimutator’ strains, are more rarely observed, particularly when the genetic change is gene inactivation. Here we identify genes that normally increase mutation rate, whose disruption gives an antimutator phenotype. Given the importance of mutation rates in determining adaptation to new environments (Swings et al. 2017), such genes could provide useful targets for ‘anti-evolution’ drugs (Cirz et al. 2005; Alam et al. 2016; Ragheb et al. 2019; Carvajal-Garcia et al. 2024).

Most previous research into antimutators focuses on secondary mutations that reverse a preexisting mutator phenotype towards the wildtype phenotype (e.g. (Fijalkowska et al. 1993; Schaaper 1993; Schaaper 1996; Wielgoss et al. 2012; Swings et al. 2017; Sprouffske et al. 2018; Ho et al. 2021)). Though rare, antimutator alleles in a wildtype background have been identified: *mfd* knockout in multiple bacterial species (Ragheb et al. 2019); mutants of *dnaE* in *Escherichia coli* (Oller and Schaaper 1994); various mutations of DNA polymerase (gene 43) of phage T4 (Drake 1993); and a temperature sensitive mutant of *purB* in *E. coli* (Geiger and Speyer 1977; but see Schaaper and Dunn 2001). These genes are integral to DNA replication and repair pathways, indicating that other components of these pathways might also have the potential to act as antimutators, although they have been less frequently identified or studied.

One under-explored gene family in this context encode the Nudix hydrolase proteins. The Nudix hydrolases were originally defined by their ‘house-cleaning’ role in degrading potentially deleterious metabolites (Bessman et al. 1996). Specifically, they were characterised by their activity in hydrolyzing nucleoside diphosphate derivatives, though substrates outside of this group have also been identified (Bessman et al. 1996; McLennan 2005; Bessman 2019). The only enzyme from this family to have been well studied in relation to mutational dynamics is MutT (also known as NudA). The nudix knockout Δ*mutT* produces a mutator phenotype in bacteria (Maki and Sekiguchi 1992), yeast (homolog *PCD1*) (Nunoshiba 2004; Krašovec et al. 2017) and human cancer (homolog *MTH1*) (Tsuzuki et al. 2001). As well as increasing mutation rates, Δ*mutT* has been shown to alter the mutational spectrum via an increase in the relative rate of A>C transversions (Tajiri et al. 1995). MutT is also involved in density associated mutation rate plasticity, the widely conserved negative association between microbial mutation rates and final population density in batch culture (Krašovec et al. 2017). While wildtype strains of *E. coli* and *S. cerevisiae* show the expected association between population density and mutation rate, deletants of *mutT* and *PCD1,* respectively, show a constant, high, mutation rate across a range of final population densities (Krašovec et al. 2017; Krašovec et al. 2018). The role of MutT in density associated mutation rate plasticity (DAMP) underscores its importance in controlling mutation rate dynamics. In contrast to *mutT*, little is known about the mutational consequences of knocking out other members of the nudix hydrolase family (Kamiya et al. 2003; Hori et al. 2006; McLennan 2012).

We begin this study by examining mutation rates across a set of Nudix hydrolase knockouts in *E. coli*. We find that 4 of the 10 knockouts show a significant antimutator phenotype, indicating that their gene products contribute to mutagenesis. We characterise the mutational spectrum of one of these newly identified antimutators, Δ*nudJ,* identifying a significant shift towards A>C transversions. Interestingly, unlike the mutator Δ*mutT*, which also increases the proportion of A>C transversions, the Δ*nudJ* antimutator achieves this shift by selectively decreasing the rate of A>G transitions. Following recent work in this area (Couce et al. 2013; Cano et al. 2022; Sane et al. 2023; Tuffaha et al. 2023) we explore whether the altered mutational spectrum of Δ*nudJ* results in differing fitness outcomes following selection for resistance to the antibiotic rifampicin. We find that the A>C dominated spectrum of Δ*nudJ* results in a bias towards less fit mutations. Therefore Δ*nudJ* adaptation to rifampicin is impaired both by a reduction in the total mutation rate and a bias towards high cost mutations.

## Results

### Several Nudix hydrolase knockouts show an antimutator phenotype

The Nudix hydrolase MutT plays a vital role in maintaining the wildtype mutation rate in *E. coli* (Maki and Sekiguchi 1992). However, *E. coli* contains a further 13 nudix hydrolases, whose effects on mutation rate have, in most cases, not been reported (though see Kamiya et al. 2003; Hori et al. 2006). We estimated mutation rates across single-gene deletants of a further ten Nudix hydrolases (out of 14 known *E. coli* proteins with a nudix domain) to systematically assess the roles of this enzyme family in mutagenesis (Figure 1). The magnitude of mutation rate change in Δ*mutT* strains (estimated to be between 50-fold (Swings et al. 2017) and 139-fold (Sprouffske et al. 2018) higher than the wildtype) enabled its early discovery as a mutator (Treffers et al. 1954). In contrast, the Nudix knockouts surveyed here show much more subtle effects on mutation rate; none show significant increases in mutation rate, but 4 of the 10 tested (Δ*nudB*, Δ*nudF*, Δ*nudH* and Δ*nudJ*) show significant reductions in mutation rate, between 1.8 and 4.5 fold lower than the wildtype (Figure 1, Table S1). It is important to note that our estimation of spontaneous mutation rate is based on rifampicin resistance which is due to single nucleotide polymorphisms (SNPs) and small indels; it does not consider larger indels, local copy number alterations or chromosomal rearrangements (Garibyan 2003).

**Figure 1:**
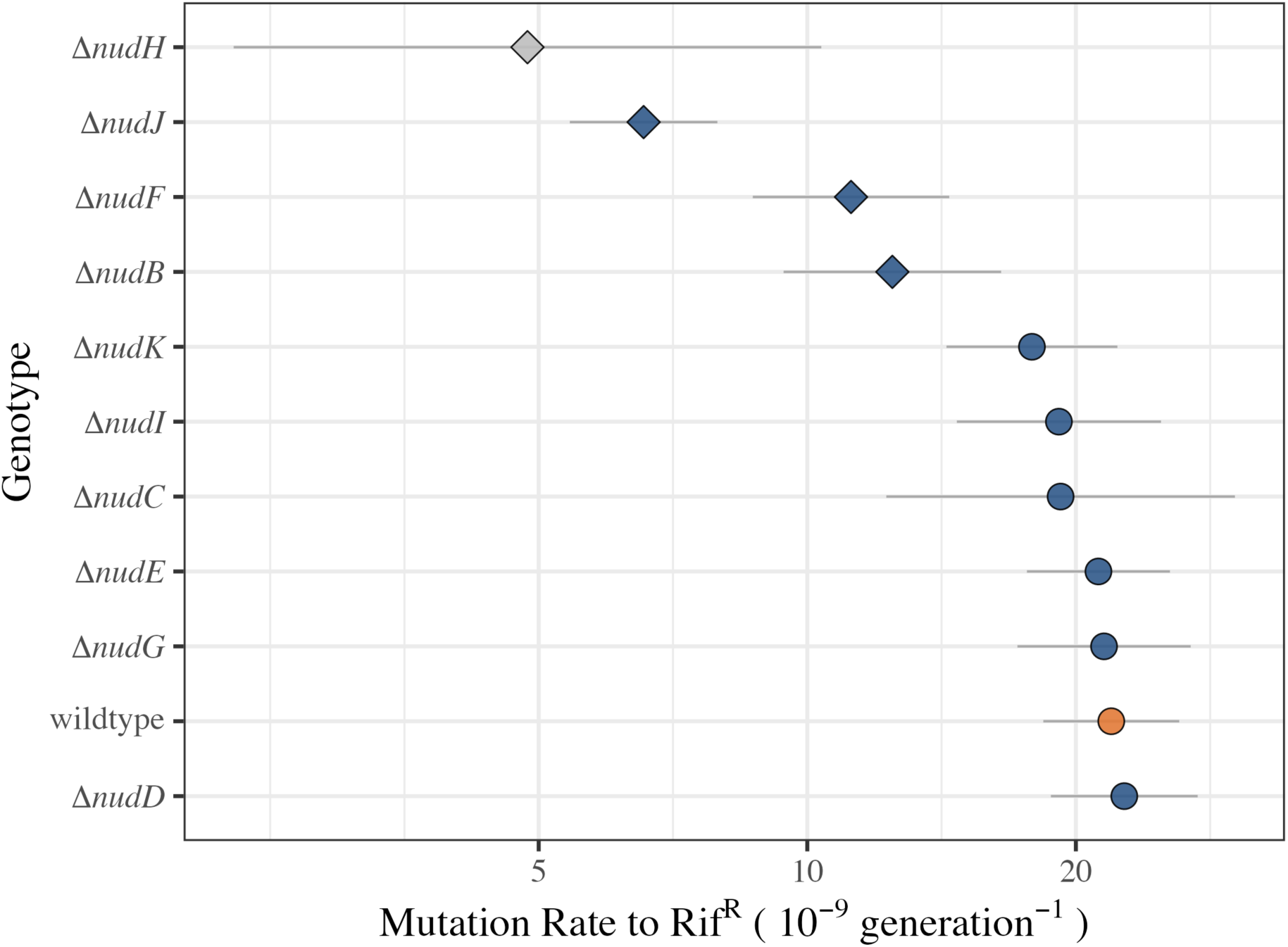
Mutation rate in Nudix hydrolase knockouts. Points show mutation rate per genome per generation and errorbars show 95% confidence intervals. Diamonds indicate strains with a significantly different mutation rate to the wildtype BW25113 see Table S1. The wildtype BW25113 is highlighted in orange. Δ*nudH* is shown in grey as this point is an extrapolation (see text). Each of these estimates and error derives from Statistical Model 1 fitted to multiple individual mutation rate estimates, collected at different population densities and accounting for observed variation of rate with density (see Supplementary Statistical Methods). Mutation rates shown here are estimated at a density of 1.47 × 10^8^ CFU.mL^-1^. Both the underlying data and fitted model are shown in Figure S1. Genotypes shown from top to bottom are: Δ*nudH* (*N*_Fluctuation Assays_ =12, *N*_Parallel Cultures_ =195), Δ*nudJ* (*N*_FA_ =47, *N*_PC_ =1089), Δ*nudF* (*N*_FA_ =16, *N*_PC_ =259), Δ*nudB* (*N*_FA_ =14, *N*_PC_ =228), Δ*nudK* (*N*_FA_ =14, *N*_PC_ =226), Δ*nudC* (*N*_FA_ =12, *N*_PC_ =193), Δ*nudI* (*N*_FA_ =14, *N*_PC_ =225), Δ*nudG* (*N*_FA_ =14, *N*_PC_ =226), Δ*nudE* (*N*_FA_ =14, *N*_PC_ =228), wildtype BW25113 (*N*_FA_ =92, *N*_PC_ =1613), Δ*nudD* (*N*_FA_ =14, *N*_PC_ =227).

The largest fold change in mutation rate shown is in Δ*nudH.* However, a growth defect in this strain prevents it from reaching comparable density to the other strains. Given the known effect of population density on mutation rates (Krašovec et al. 2017; Green et al. 2024), the mutation rate estimates shown in Figure 1 are at the mean density across all assays. The Δ*nudH* mutation rate shown is therefore an extrapolation to its mutation rate at a higher density than it ever achieves in our experiments (see Figure S1), giving the estimate large error bars and questionable value. After Δ*nudH*, the largest change is seen in Δ*nudJ,* which is a clear outlier within the remaining Nudix deletant strains tested here, showing a mutation rate reduction of 3.3-fold (Figure 1). We therefore focus primarily on Δ*nudJ* going forward.

### NudJ’s metabolic contribution to mutagenesis via prenyl and nucleotide metabolism

The observation of a strong antimutator phenotype in Δ*nudJ* indicates that some activity of NudJ is contributing to mutagenesis in *E. coli*. NudJ is known to show strong dephosphorylating activity *in vitro* against dimethylallyl-diphosphate (DMAPP), producing dimethylallyl-monophosphate (DMAP). DMAP is required for the production of the UbiD cofactor prenylated flavin mononucletide (prFMN) which is essential for ubiquinone biosynthesis (Gulmezian et al. 2007; Payne et al. 2015; Wang et al. 2018), (Figure S2). DMAPP dephosphorylation activity is seen in 3 of the 4 identified antimutators (NudB, F & J but not NudH) but only in one of the 7 non-antimutators (NudI). As well as dephosphorylating DMAPP, NudJ also dephosphorylates the essential cofactor thiamin phyrophospate (TPP) and its precursor in thiamin salvage, 4-amino-2-methyl-5-hydroxymethylpyrimidine pyrophosphate (HMP-PP), *in vitro* (Lawhorn et al. 2004), (Figure S2). Both the dephosphorylation of TPP and HMP-PP by NudJ may diminish the available pool of TPP. Interestingly, the enzyme ThiM is also involved in both DMAP and TPP synthesis. Like NudJ, ThiM can produce DMAP (using prenol rather than DMAPP as a substrate) (Wang et al. 2018). However, in the TPP salvage pathway ThiM has an opposing effect to NudJ: it phosphorylates 4-methyl-5-hydroxyethylthiazole (HET) to produce HET-P, aiding rather than antagonising the salvage of TPP (Mizote and Nakayama 1989), (Figure S2).

We therefore hypothesise that NudJ affects mutation rate either via its role in DMAP or in TPP metabolism. Because ThiM has a similar role to NudJ in one case and an antagonistic role in the other, both these hypotheses may be tested via the Δ*thiM* mutant. If what matters for mutation rate is the shared role of ThiM and NudJ in DMAP formation, we would expect a decreased mutation rate in a Δ*thiM* strain. On the other hand, if TPP metabolism is key to mutation rate, we would expect an increase of mutation rate in the Δ*thiM* strain. We found a non-significant decrease in the mutation rate of Δ*thiM* compared to the wildtype (t = -0.599, DF = 283, *P =* 0.549, Statistical Model 1) (Figure S3). It is therefore more likely that NudJ affects the mutation rate via its role in DMAPP dephosphorylation than in thiamin salvage. The lack of a significant effect of *thiM* deletion on the mutation rate is consistent either with some role of DMAPP dephosphorylation or with NudJ’s role in mutagenesis being unrelated to either its role in DMAP production or in thiamin salvage.

NudJ has also been shown to dephosphorylate all deoxy- and ribonucleotides in their tri- and dinucleotide forms with a preference for GDP (Xu et al. 2006). The role of NudJ in mutagenesis may come via altering the nucleotide pool composition rather than through it’s activity in DMAP production and thiamin salvage. We hypothesise that NudJ affects the mutation rate via elevation of the internal nucleotide pool (similar to the ‘dNTP mutator’ effect described by Gon et al. (2011)). To test this hypothesis we measured the internal ATP concentration, the most abundant nucleotide, in cultures of the wildtype and the nudix hydrolase knockouts. Δ*nudJ* shows the lowest internal ATP concentration observed across the tested strains, with an ATP concentration significantly lower than the wildtype (t_DF=33_ = -2.82, *P* = 0.00794, Statistical Model 2, Figure S4). These results are consistent the hypothesis that NudJ contributes to mutagenesis via elevation of the internal nucleotide pool.

### Epistatic outer membrane mutations reverse the Δ*nudJ* antimutator phenotype

Significant mutation rate reductions in wildtype *E. coli* are uncommon. We therefore verified our findings in the Δ*nudJ* strain by assaying mutation rates in a second independent *nudJ* deletant as well as 3 further colonies from the original deletant stock (totaling 5 strains: 4 from the original deletant and 1 from an independent deletant, all derived from the Keio collection (Baba et al. 2006)). Within this set of *nudJ* knockouts we found clear variation in mutation rates; three show the previously observed antimutator phenotype while two show a wildtype mutation rate (Figure S5). All of the strains with an antimutator phenotype derive from the original knockout and have no secondary mutations in coding regions. In contrast, the two strains showing a wildtype mutation rate carry secondary mutations in coding regions: the first, from the original knockout, has an IS1 insertion within the first quarter of the *waaZ* coding sequence; the second, deriving from an independent knockout, has both an IS5 insertion in the first half of the *csgF* coding sequence and a frameshift +CG mutation within the first 15 bases of the *ispA* coding sequence (Figure 2). It is likely that these IS and frameshift mutations are functionally equivalent to whole gene deletants. In both of these revertant strains, IS elements disrupt genes involved in the barrier between cells and their environment; WaaZ is involved in formation of the lipopolysaccharide core which forms part of the outer membrane (Frirdich et al. 2003) whilst CsgF is required for efficient assembly of curli, the major proteinaceous component of the extracellular matrix (Barnhart and Chapman 2006). Mutants in the *waaQGPSBIJYZK* operon have been shown to be deficient in curli production (Smith et al. 2017), potentially explaining the similar phenotypes of strains with each of these secondary mutations.

**Figure 2:**
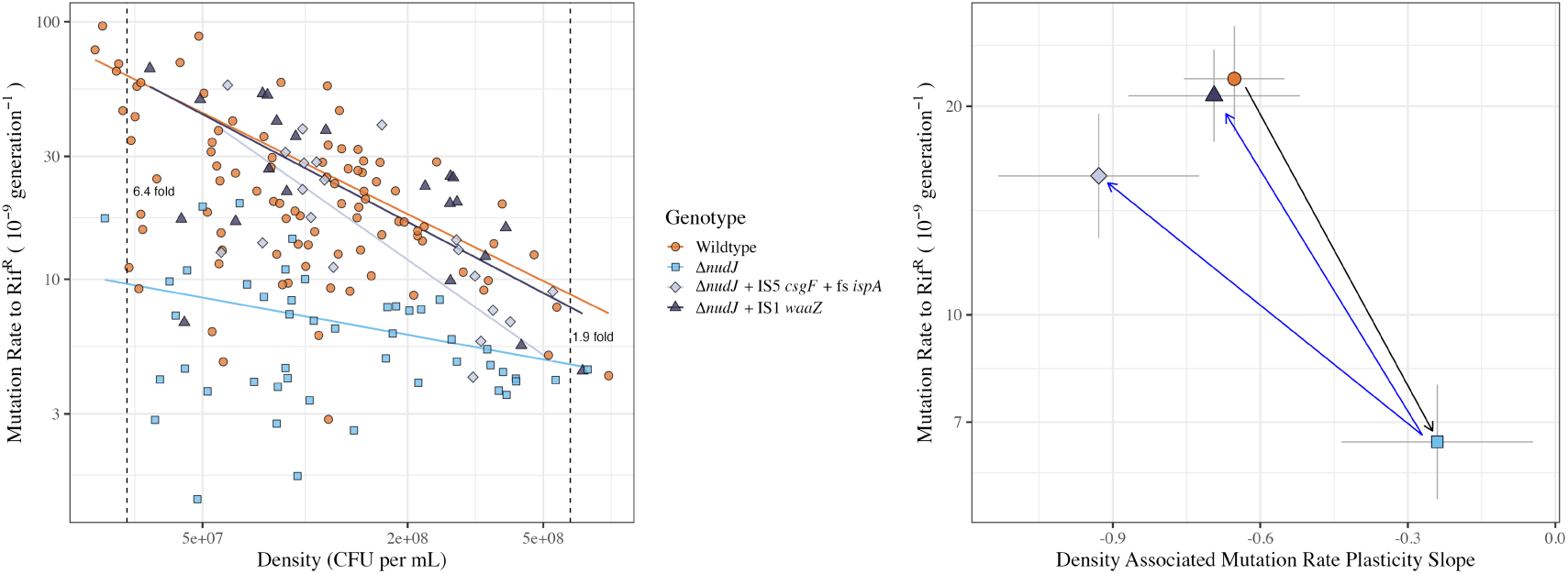
Mutation rates in wildtype BW25113 and Δ*nudJ* strains. A) Mutation rate to rifampicin resistance (mutational events per generation (x10^9^)) plotted as a function of final population density (CFU per mL). Orange circles: BW25113 wildtype (*N*_Fluctuation Assays_ = 92, *N*_Parallel Cultures_ = 1613); Pale blue squares: Δ*nudJ* (*N*_FA_ = 47, *N*_PC_ = 1089); Pale grey diamonds: Δ*nudJ* + *waaZ* IS1(*N*_FA_ = 22, *N*_PC_ = 478); Dark grey triangles: Δ*nudJ* + *csgF* IS5 + *ispA* FS (*N*_FA_ = 20, *N*_PC_ = 321). Dashed lines indicate population densities of 3×10^7^ and 6×10^8^ at which the fold difference in mutation rate between the wildtype BW25113 and Δ*nudJ* is estimated to be 6.4 fold (95% CI: 4.3 - 9.72) and 1.9 fold (95% CI: 1.4 - 2.5) respectively. B) Mutation rate (intercept at a density of 1.47 × 10^8^) and slope (density associated mutation rate plasticity) of lines of the fitted regressions shown in figure 2A with 95% confidence intervals (Statistical Model 1). Black arrow indicates the change in phenotype caused by *nudJ* deletion and blue arrows indicate the reversion in phenotype caused by secondary mutations. Mutation rates for wildtype and Δ*nudJ* are as in figure 1.

These secondary mutations may cause an increase in mutation rate independent of the *nudJ* allele, balancing the antimutator effect of *nudJ* deletion. Alternatively, these mutations might have no effect on their own, but epistatically interact with the *nudJ* deletion to revert the mutation rate. To test this, we assayed the mutation rate of a strain with *waaZ* deleted in a wildtype background. We focus on the *waaZ* mutant rather than the *csgF* + *ispA* mutant due to the conditional essentially of *ispA* (Fujisaki et al. 2005; Baba et al. 2006)*. waaZ* deletion failed to detectably increase the mutation rate (t _DF=18_ = -1.45, *P* = 0.919, Figure S6, Statistical Model 3). Thus, *waaZ* disruption has an epistatic effect with *nudJ* deletion, significantly elevating the mutation rate in the Δ*nudJ* background but not the wildtype background (t = 11.7 _DF=102_, *P* = 4.41 × 10^−26^, Figure 2, Statistical Model 1).

To further test the hypothesis that NudJ modifies mutation rates via elevation of the internal nucleotide pool we also measured internal ATP in the revertant Δ*nudJ* strains (Figure S4). We find that, unlike Δ*nudJ* alone, Δ*nudJ* + IS1 *waaZ* does not significantly differ from the wildtype (t_DF=33_ = -0.0155, *P* = 0.988, Statistical Model 2) whilst Δ*nudJ* + IS5 *csgF* + fs *ispA* actually increases the internal ATP concentration relative to the wildtype (t_DF=33_ = 3.47, *P* = 0.00191, Statistical Model 2), contrasting with the decrease in ATP pool seen in the original Δ*nudJ* strain. This supports our hypothesis that the mutation rate decrease seen in Δ*nudJ* is caused by a change in the internal nucleotide pool, as indicated by a depletion of ATP which is recovered to wildtype concentrations, or above, in the revertant strains.

A possible confounding factor in our discovery of Δ*nudJ* as an antimutator is that this could simply reflect the known association between growth rate and mutation rate (Novick and Szilard 1950; Chu et al. 2018; Maharjan and Ferenci 2018; Liu and Zhang 2019). Previous work has generally identified a negative correlation between microbial mutation rates per generation and growth rate (Novick and Szilard 1950; Maharjan and Ferenci 2018; Liu and Zhang 2019). We would therefore expect that if growth rate is driving the antimutator phenotype of Δ*nudJ*, this strain will have an increased growth rate. Instead, we see no change in the growth rate of the Δ*nudJ* strain (Figure S7) although the carrying capacity is reduced in this strain (Figure S7 and Figure S8). This reduction in carrying capacity is not restored by the secondary mutations which restore the wildtype mutation rate, again indicating that the role of NudJ in mutation rate determination is not due to changes in growth dynamics (Figure S8).

### The antimutator phenotype of Δ*nudJ* is environment-dependent

Mutation rate is known to be negatively correlated with population density in wildtype *E. coli* (Krašovec et al. 2014). In addition to showing a reduced mutation rate, Δ*nudJ* also shows a significantly weaker relationship between density and mutation rate compared to the wildtype (Δ*nudJ* slope = -0.23±0.19, wt slope = -0.65±0.11, 95% CI, t _DF=102_ = 4.17, *P* = 1.24 × 10^−4^, Statistical Model 1). Because of this change in environmental responsiveness, the antimutator phenotype of Δ*nudJ* is strongest at low densities. For example at a low population density of 3×10^7^ CFU mL^-1^ we see a mutation rate decrease of over 6-fold in Δ*nudJ* as compared to the wildtype, in contrast at a higher density of 6×10^8^ CFU mL^-1^ Δ*nudJ* only shows a 1.9-fold reduction in mutation rate (Figure 2). Density associated mutation rate plasticity (DAMP) is largely driven by elevation in the rate of A>G mutations at low density (Gifford et al. 2024); we therefore hypothesised that Δ*nudJ* may be less prone to this class of SNP.

### Mutational spectrum is depleted in A>G transitions in antimutator Δ*nudJ*

To test for changes in mutational spectrum in the Δ*nudJ* background we collected spontaneously occurring rifampicin resistant mutants from mutation rate assays with the wildtype, Δ*nudJ* and Δ*nudJ* + IS1 *waaZ* strains. These mutants were collected from low density populations (defined as those grown in 80mg.L^-1^ glucose) as it is in this condition that we see the largest difference in mutation rate and therefore might expect to see the greatest deviation in spectra between Δ*nudJ* and the wildtype. We sequenced the rifampicin resistance determining region (RRDR) of *rpoB* in resistant strains with each of these three genetic backgrounds. Although the distribution of mutational classes observed within the RRDR does not reflect that across the whole genome, this method does allow for robust comparison of the spectra of point mutations among strains (Garibyan 2003).

Δ*nudJ* shows a significantly different mutational spectrum to the wildtype ancestor (LR _DF=7_ = 17.6, *P* = 0.0279, Statistical Model 4). However, the secondary mutation in Δ*nudJ* + IS1 *waaZ* causes the mutational spectrum to revert to that of the wildtype (LR _DF=7_ = 5.2, *P* = 0.6309216, Statistical Model 4) (Figure 3). The most striking differences in the spectrum introduced by Δ*nudJ* are a decrease of over 50% in the proportion of A>G transitions (*χ*^2^ = 7.62, DF = 1, *P =* 0.0231), and a concurrent increase in A>C transversions (*χ*^2^ = 10.7, DF = 1, *P =* 0.00853) (Table S3). This is consistent with our prediction that a reduction in A>G transitions is driving the observed reduction in DAMP (Gifford et al. 2024) (Figure 2). While the relative shifts in the mutational spectrum of Δ*nudJ* potentially account for its loss of DAMP, it is important to note that the absolute rates of all mutational classes, including A>C transversions, are decreased to differing degrees in Δ*nudJ* (Figure S9). The elevated rate of A>G transitions seen here in the presence of NudJ has also been observed under thymine starvation where this phenotype has been suggested to result from imbalances in the intracellular nucleotide pool (Kunz and Glickman 1985). This is consistent with our hypothesis that the role of NudJ in mutagenesis is driven by its nucleotide dephosphorylase activity, altering the intracellular nucleotide pool composition (Xu et al. 2006).

**Figure 3:**
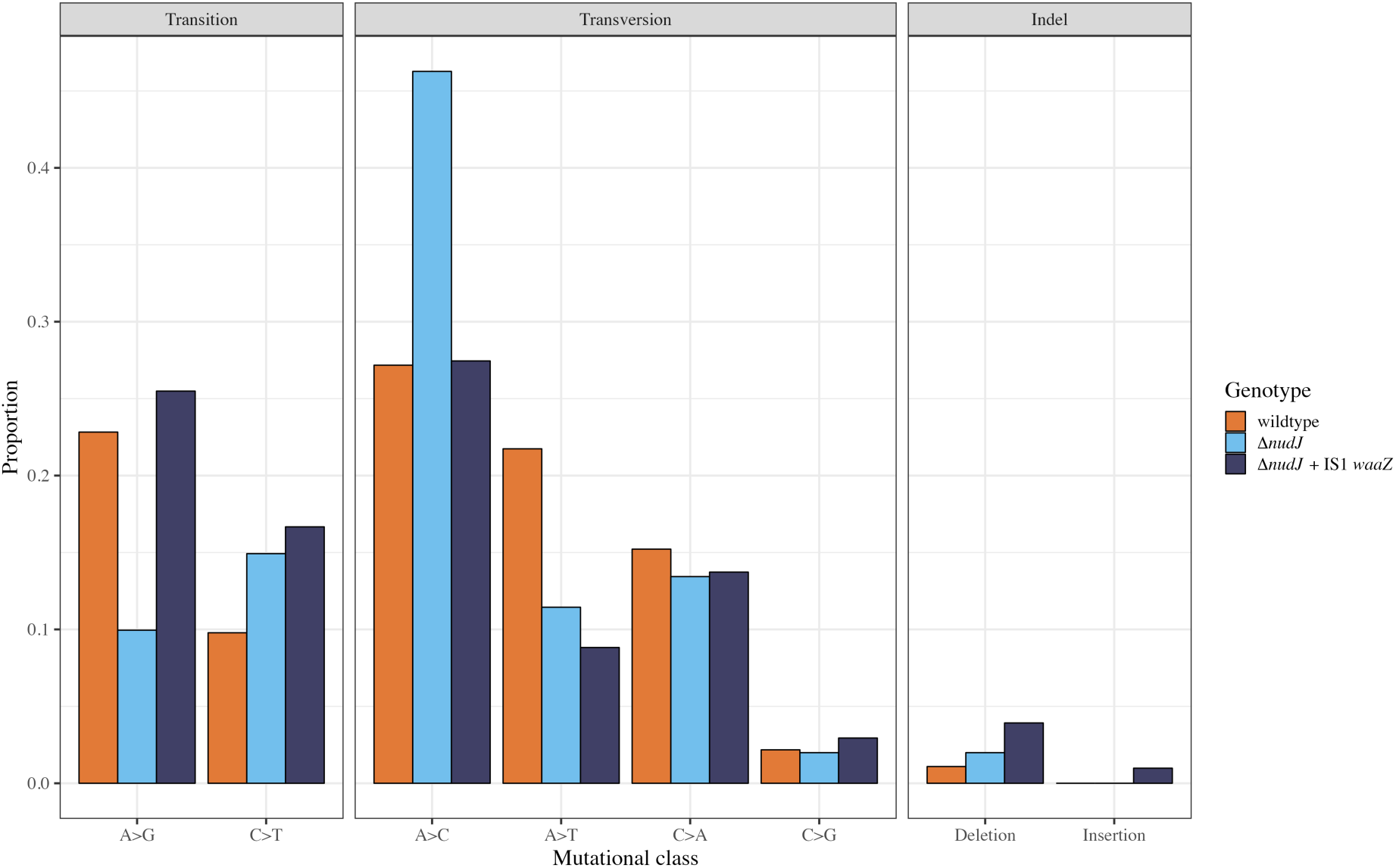
Mutational spectra of rifampicin resistance (rifR) mutations in wildtype BW25113, Δ*nudJ* and Δ*nudJ* + *waaZ* IS1 genetic backgrounds. Proportion of identified rifR mutations in the RRDR from each mutational class are shown for wildtype BW25113 in orange (*N* = 92), Δ*nudJ* in pale blue (*N* = 201) and Δ*nudJ* + *waaZ* IS1 in dark blue (*N* = 102). All mutations are listed in Table S2.

One possible alternative explanation for the reduced muation rate of Δ*nudJ* to rifR as compared to the wildtype is that a smaller possible pool of amino acid substitutions provide rifampicin resistance in the Δ*nudJ* background. Using the observed set of sequenced rifR mutants from these genetic backgrounds we can test this alternative hypothesis. Across 92 mutants in the wildtype background, 30 unique AA substitutions were observed whilst in the Δ*nudJ* background 39 AA substitutions were observed across 201 mutants (Table S2, Figure S10). If we randomly subset the Δ*nudJ* mutants to a sample of 92 we observe a mean average of 30.2±4.1 unique AA substitutions (95% prediction interval, 1×10^6^ repeats), not significantly different from the 30 unique substitutions seen in the wildtype. This suggests that the possible pool of amino acid substitutions providing rifampicin resistance does not differ in size between the wildtype and Δ*nudJ* backgrounds.

### Mutational hotspots in *rpoB* do not differ between the Δ*nudJ* antimutator and the wildtype

RifR mutations are not only biased by mutational class but also by genomic location, indicating the presence of mutational hot and cold spots within the sequenced region of *rpoB* (Figure S11) (as previously observed by (Wolff et al. 2004)). For example, while we find A>C transversions at position 1687 (T563P) to account for 63/139 observed A>C mutations observed across all strains, the least frequent A>C transversion, at position 1598 (L533R) was observed only 3 times. We find a significant effect of position on mutant frequency, accounting for differences in the relative rates of mutational classes, indicating that hotspots exist in the rifampicin resistance determining region of *rpoB* (Deviance_DF=48_ 88.0, *P* = 3.76 × 10^−4^). This is a conservative estimate given the likely existence of unobserved rifR mutants (Sun and Lind 2023). These hotspots do not correlate with fitness of the resultant mutant, discounting explanations of biased mutant selection or an evolutionary bias towards low cost resistance mutations (Figure S12); independence of mutational frequency and mutant fitness has also been previously reported in *Pseudomonas aeruginosa* (MacLean et al. 2010).

It is possible that these hotspots, like the mutational spectra, will differ between genetic backgrounds. This would indicate that not only does *nudJ* deletion bias *E. coli* towards different mutational classes but towards different specific mutations within these classes. We find no significant evidence for changes in the strengths of mutational hotspots within any mutational class between the wildtype, Δ*nudJ* and Δ*nudJ* + IS1 *waaZ* (Deviance_DF=45_ = 18.0, *P* = 0.9999, Statistical Model 5 Figure S13).

### The available distribution of growth effects is altered by mutational spectrum

Because of the altered mutational spectrum of Δ*nudJ*, the probabilities of each particular spontaneous resistance mutation arising, i.e. mutation accessibility, is changed. For example, 12% of all rifR mutants identified in the wildtype background were D516G, caused by an A>G transition. In contrast, only 5% of rifR mutants in the Δ*nudJ* background were accounted for by this DNA and amino acid (AA) substitution. Particularly with rifampicin, where the target of resistance, *rpoB*, is an essential gene, different resistance mutations have different fitness costs associated with them (MacLean et al. 2010; Maharjan and Ferenci 2017). We therefore hypothesised that the distribution of fitness effects (DFE) of accessed rifR mutants (termed the DFE_*β*_ by (Tuffaha et al. 2023)) will differ depending on the genetic background in which they evolved. The DFE_*β*_ can be compared to the null expectation that each mutant will be observed an equal number of times, we will refer to this distribution as the DFE_all_ as in (Tuffaha et al. 2023).

In order to characterise the DFE_*β*_ for each of our strains we measured the growth effects (as the area under the curve (AUC) of 24 hr growth curves) in the common MG1655 background of 27 RpoB amino acid substitutions. This type of growth assay is a good proxy for competitive fitness by Vogwill and MacLean (2014). The 27 amino acid substitutions selected account for 83% (326/395) of our mutants with identified mutations (the remaining 18.5% being 29 rare mutations). Given the environmental dependence of the fitness effects of *rpoB* mutations (Hall 2013; Gifford et al. 2016; Soley et al. 2023), we assessed mutant growth across 12 different environments to achieve a more general characterisation of the impact of mutational spectrum on the DFE_*β*_. Our environments include all possible combinations of temperature (25℃ vs 37℃ vs 42℃), nutrient (minimal M9 vs rich LB) and rifampicin (presence vs absence).

To test the hypothesis that mutants expected to evolve in different backgrounds will have different fitness distributions, we weighted the growth measured for the *rpoB* mutants in the wildtype MG1655 background according to the estimated probability of observing the underlying DNA mutation in each strain’s mutational spectrum and compared those strains (see Statistical Model 6). We find that mutants evolved in the Δ*nudJ* background have a significantly lower average fitness than those evolved in the wildtype background (t_DF=1572_ = - 4.26, *P* = 2.12 × 10^−5^, Statistical Model 6) (Figure 4, Figure S14). Therefore, not only is rifampicin resistance observed less frequently in Δ*nudJ*, due to the reduced mutation rate, but when it does arise, greater fitness costs are typically incurred. We can also estimate the fitness costs of rifR from the data going into our mutation rate estimates. These are counts of rifR colony forming units, which will be reduced if resistance mutations carry a cost in the non-selective environment of the fluctuation assay. By co-estimating the average mutant fitness with the mutation rate, we see that rifR mutants do indeed show a lower average fitness in the Δ*nudJ* background relative to the wildtype (Figure S15).

**Figure 4:**
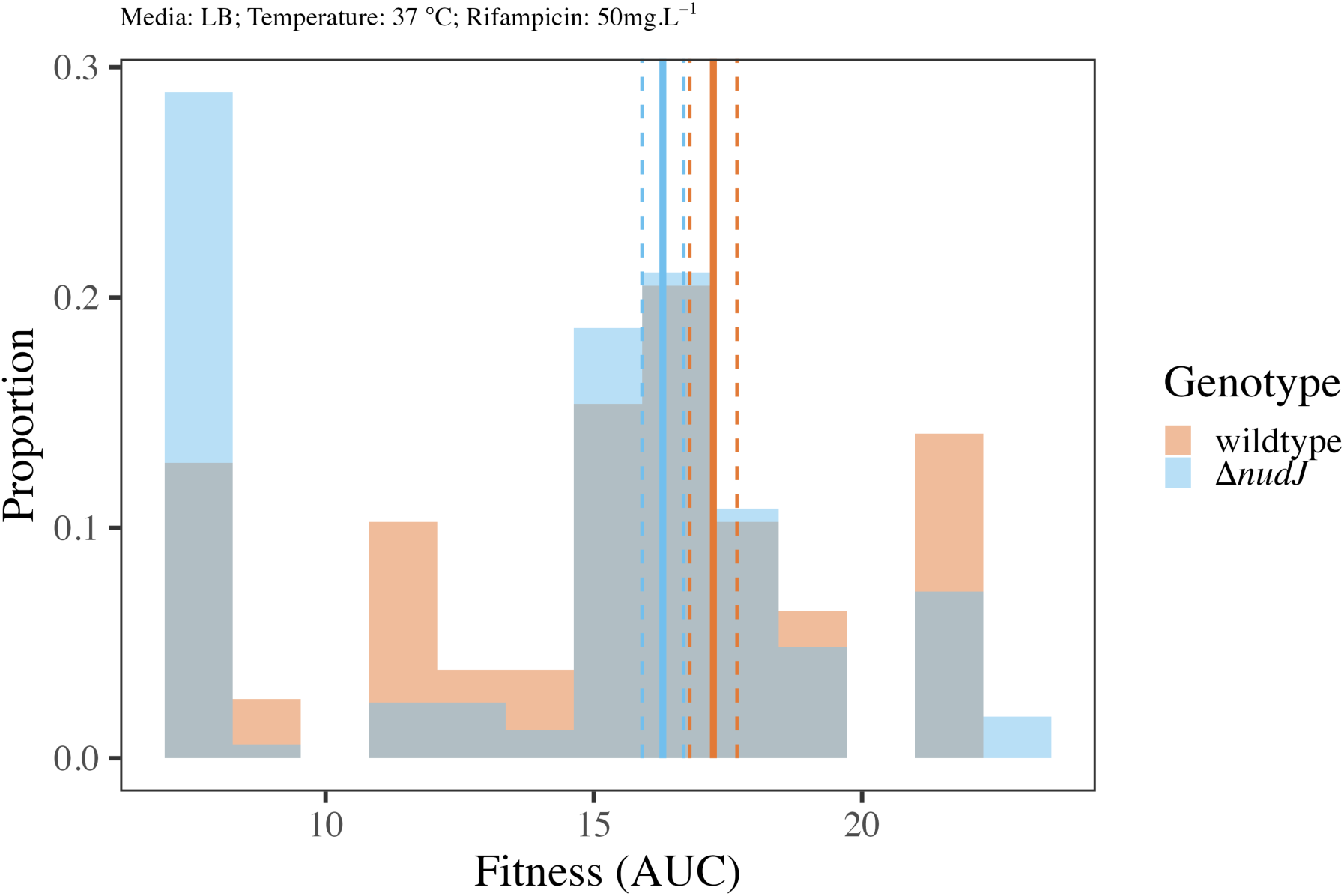
The fitness of evolved rifR mutants depends on mutational spectrum. Histogram of growth for observed mutants evolved in the Δ*nudJ* vs wt background. Fitness for each mutant is measured in the common MG1655 background grown in rich LB media with rifampicin. Data for all other environments is shown in Figure S14. Mean and 95% CI for fitness of mutants evolved in each genetic background shown by solid and dashed lines as predicted from Statistical Model 6.

Theory developed by Tuffaha et al. (2023) describes how the effect of mutation rate, the beneficial fraction of the DFE_*β*_, and the average magnitude of beneficial mutations combine multiplicatively to determine the chances of a beneficial mutation carrying a strain to fixation. Given a 6.4-fold difference in mutation rate, one would expect that in a population with equal numbers of wildtype and Δ*nudJ* individuals, the wildtype would be 6.4 times more likely than Δ*nudJ* to be carried to fixation by a beneficial *rpoB* mutation. However once the fitness effects of rifR mutations observed in each genetic background are also accounted for, this ratio of fixation probabilities can be as high as 11.7, varying by the environment in which the DFE is measured (Table S4). This calculation is a simplification, not least because it considers only the mutation rate and spectra differences observed in low density populations grown in minimal media (Figure 3). However, it is a demonstration of the substantial role that the mutational spectrum can play in the evolutionary fate of the resistant strains evolved in antimutators. This effect of mutational spectrum is especially pronounced given the relatively modest fold changes in mutation rate seen here with an antimutator as compared to the much larger fold changes seen in mutator strains (Lee et al. 2012; Foster et al. 2015) for which the evolutionary effects of mutational spectrum changes have been previously considered (Couce et al. 2013).

Our finding that Δ*nudJ* preferentially accesses rifampicin resistant mutants incurring greater fitness costs than the wildtype, may or may not be generalisable to other antibiotics. Hypothetically, during adaptation to another antibiotic Δ*nudJ* may access mutants incurring the same fitness costs as wildtype or it may access mutants with higher or lower fitness costs than the wildtype. We assessed the effect of the Δ*nudJ* mutational spectrum on the fitness and growth effects of resistance mutations to the antibiotics nalidixic acid and trimethoprim using data from (Harmand et al. 2017) and (Palmer et al. 2015) respectively. We used a bootstrapping approach to sample mutants under the spectra of Δ*nudJ* and the wildtype independently before comparing the fitness effects of mutants in these two pools (see Methods section ‘Bootstrapping fitness differences…’). In the case of nalidixic acid we find a consistently negative effect of the Δ*nudJ* mutational spectrum on the average fitness of accessible mutants, consistent with our results for rifampicin Figure S16. This effect is seen when fitness of the mutants is measured either in the absence of antibiotics or in the presence of a lethal contentration of nalidixic acid. In contrast, the difference in the average growth of trimethoprim resistant mutants accessed by Δ*nudJ* vs the wildtype depends on the environment in which growth is measured. Mutants accessed by Δ*nudJ* show consistently poorer growth that those accessed by the wildtype in sub-inhibitory concentrations of trimethoprim. However, in higher concentrations of trimethoprim this effect is reversed and the mutants accessed by Δ*nudJ* have greater growth than those accessed by the wildtype. These findings demonstrate that rifampicin is not unique in the Δ*nudJ* strain preferentially accessing low fitness resistance mutants. Nonetheless, the impact of mutational spectra on the fitness effects of resistance mutations does depend on the antibiotic in question, and its interaction with the environment in which fitness is measured.

### RifR mutant fitness is correlated with the destabilising effect of mutations

To test the basis for the variation among the different DFE_*β*_, we asked what explanatory factor could be determining the fitness of individual *rpoB* mutants. One potential cause of mutant fitness variation is the change in protein stability caused by the change to the AA sequence. Previous evidence suggests that mutations with more destabilising effects on the protein will impair function, and therefore fitness (Bershtein et al. 2012; Sarkisyan et al. 2016). Using an empirical forcefield implemented by FoldX (Schymkowitz et al. 2005) we estimated the change in the Gibbs free energy of folding (ΔΔG) caused by each of the observed mutations. For each mutant we estimated the ΔΔG based on the structure of the RpoB subunit alone and of the whole RNA polymerase (RNAP) complex. These estimates were correlated, though with outliers corresponding to one specific AA position (overall Spearman’s rho = 0.558, *P* = 0.00291, Figure S17). In both cases the majority of mutations were predicted to be destabilising (ΔΔG>0) (Table S2). When fitting a statistical model to predict fitness, as measured by the proxy of AUC, from ΔΔG of the RpoB subunit and ΔΔG of the RNAP complex we find that only the former effect is significant. We find a significant negative correlation between AUC and ΔΔG of the RpoB subunit (F = 18.9, DF = 1, 318, *P* = 1.84 × 10^−5^, Statistical Model 7, Figure S18), consistent with the hypothesis that more destabilising mutations will impair fitness to a greater extent. However, although this correlation is significant, the proportion of observed variance in AUC explained by the ΔΔG of the RpoB subunit is small (see Statistical Model 7). These results indicate that protein stability, while significant, is not the sole determinant of the pleiotropic growth effects of *rpoB* mutations. Therefore, although a bias towards selecting mutations with more destabilising effects on protein folding may lead to negative shifts in the DFE_*β*_, much of the variance in fitness effects of mutations is not explained by protein stability.

## Discussion

By characterising the roles of ten Nudix hydrolases in spontaneous mutagenesis we have highlighted disruption of NudJ, NudB, NudF and NudH as novel pathways to reduced mutation rates in *E. coli* (Figure 1). We focused on the *nudJ* deletant, finding that not only is the mutation rate reduced, but the mutational spectrum is also significantly altered in this strain (Figure 3). Δ*nudJ* has a mutational spectrum significantly depleted in A>G transitions and instead has a much higher proportion of A>C transversions than the wildtype. This change in spectrum causes Δ*nudJ* to preferentially access rifampicin resistance mutations of significantly lower fitness than those accessed by the wildtype (Figure 4, Figure S14), with low fitness being associated with a predicted destabilising effect of the resistance mutations on the RNA polymerase subunit (Figure S18). Both the antimutator phenotype and altered mutational spectrum of Δ*nudJ* are reversed by a secondary mutation in *waaZ*, demonstrating the complexities underlying the genetic determination of microbial mutation rates (Figure 2, Figure 3). Taking Δ*nudJ* as a model we show how multiple factors influencing evolvability (mutation rate, mutation rate plasticity and shifts in the available DFE caused by mutational spectrum) can be simultaneously altered by a single gene knockout. This study highlights the importance of considering not only the mutation rate but also the mutational spectrum and environmental responsiveness in these traits if we are to understand the evolutionary impacts of mutation rate modifying alleles. If leveraged well, these additional properties of antimutators will enhance our ability to control evolutionary trajectories.

Although we find significant evidence of mutational hotspots in *rpoB* (Figure S11), we do not find the location of these hotspots to be altered in the Δ*nudJ* antimutator strain (Figure S13). This echoes the findings of Shepherd et al. (2022) who find that a transversion based hotspot identified in wildtype *Pseudomonas fluorescens* remains active in a transition biased Δ*mutS* mutator. These findings indicate that, while changes in mutational spectra and mutation rates are commonly correlated due to a shared cause (e.g. Lee et al. 2012; Garushyants et al. 2024), in this case most likely associated with the nucleotide pool, mutational hotspots are controlled by distinct mechanisms (reviewed by Horton and Taylor 2023). While mutation rates and spectra are typically controlled by the genome wide effects of mutation rate modifying alleles, mutational hotspots are typically determined by local sequence context. For example, the most frequently observed mutation in this study (an A>C mutation at position 1687 leading to AA substitution T563P) alters the sequence at positions 1685-1691 from ‘CG-AA**<u>A</u>**CCCC-TG’ to ‘CG-AA**<u>C</u>**CCCC-TG’; this is consistent with the mutagenic effect of G runs 5’ of a T nucleotide (as would be seen on the opposite, leading, strand, reading ‘CA-GGGG**<u>T</u>**TT-CG’) (Cherry 2023).

This study highlights the interplay between mutation rate and spectra in determining the evolvability of a population. In the case of Δ*nudJ*, reduction of the mutation rate and a bias toward mutants of lower fitness compound to reduce evolvability to rifampicin-containing environments. It is entirely plausible that, if we had considered another drug, the mutational spectrum of Δ*nudJ* may instead have provided access to mutants with greater average fitness than the wildtype spectrum. Using bootstrap sampling we show that the mutational spectrum of Δ*nudJ* will facilitate access to nalidixic acid resistance mutations with lower fitness than those accessed by the wildtype Figure 4, Figure S14, Figure S16. In contrast, the relative growth effects of trimethoprim resistance mutations accessed under the mutational spectra of Δ*nudJ* and the wildtype vary by the environment in which growth is measured Figure S16. We can also consider previous work by Couce et al. (2013) who show that a Δ*mutT* mutator is biased towards higher cost mutations to rifampicin resistance but towards less costly mutations to streptomycin resistance when compared to the C>A biased Δ*mutY* mutator. Given the A>C dominated mutational spectrum of both Δ*mutT* and Δ*nudJ*, we could infer that a NudJ inhibitor would be more effective in preventing rifampicin resistance compared to streptomycin resistance.

The concept of antievolution drugs, capable of slowing the evolution of resistance during antibiotic treatment, sufficient that an infection is expunged either by the antibiotic or immune system before resistance arrises, has also been explored via the inhibition of RecA, which promotes antibiotic induced mutagenesis. Its inhibition can potentiate the activity of multiple classes of antibiotics (Alam et al. 2016). Bacterial populations can also evolve antimicrobial resistance through horizontal gene transfer which is simillarly potentially susceptible to anti-evolution strategies: chemically inhibiting bacterial competence can reduce the spread of antimicrobial resistance genes within an infectious population (Domenech et al. 2020).

The mutation supply rate in a population is the product of the per-genome mutation rate and population size (de Visser et al. 1999), making reduced mutation rates likely to be more consequential in small populations in which mutational supply is already more limited. Where mutation supply, and hence clonal interference, is limited, adaptation is likely to proceed by the ‘survival of the first arrival’ (MacLean et al. 2010; Cano et al. 2022). Although strongest under low mutational supply it is worth noting that mutational biases can still impact evolutionary paths under a relatively high mutation supply rate (Yampolsky and Stoltzfus 2001; Cano et al. 2022). Both mutation rate and mutational spectrum can therefore play important roles in the evolvability of a population with this effect depending on demographic factors. The possible contribution of mutations to adaptation depends on their phenotypic effects, which can vary between mutational classes (e.g. (Couce and Tenaillon 2019; Sane et al. 2023; Sane et al. 2024)). Given that mutation rate changes often come with shifts in mutational spectrum (Foster et al. 2015; Garushyants et al. 2024), there is potential for synergism or antagonism between the effects of mutation rate and mutational spectrum on the evolvability of a population. In the case of of the Δ*nudJ* strain we see a synergistic effect where both mutation rate and spectrum limit the potential for adaptive evolution to a rifampicin-containing environment.

The rarity of antimutator alleles is predicted by the ‘drift-barrier hypothesis’ which proposes that mutation rates are generally under directional selection towards a minimum and only prevented from reaching this minimum by the effect of genetic drift (Sung et al. 2012; Liu and Zhang 2021). The large effective population size of *E. coli* would suggest that most easily accessible alleles, such as those causing single gene disruption, that reduce mutation rates, should already be fixed. However, the assumption that antimutators should be selected for ignores fitness effects; for example mutation rate modifiers may have direct pleiotropic fitness consequences (Saint-Ruf et al. 2007; Fitzsimmons et al. 2018) and the fitness consequences of the particular mutations they modify may vary (Couce et al. 2013; Sane et al. 2023; Sane et al. 2024). A second important consideration is the known environment-by-genotype interactions which determine microbial mutation rates (e.g. Fowler et al. 1994; Green et al. 2024). Because of the environmental effect, an allele that appears to elevate mutation rates in the lab may not do so in the natural habitat of the organism and may therefore persist in the wild population. Therefore, the identification of antimutators under lab conditions does not itself refute the drift-barrier hypothesis and the scope for identification of as yet undiscovered antimutators may be greater than previously assumed.

Similar to Δ*nudJ*, deletion of either *mfd* in multiple bacterial species or *PSP2* in *S. cerevisiae* have previously been shown to reduce mutation rates (Ragheb et al. 2019; Liu and Zhang 2021). Mfd interacts with RNA polymerase (RNAP) to remove blocked RNAP and recruit repair enzymes to DNA lesions as well as aiding transcription of hard to transcribe regions (Selby et al. 1991; Ragheb et al. 2020), while PSP2 binds to specific mRNAs, promoting their translation (Liu and Zhang 2021). Both of these proteins are involved in RNA transcription or translation whereas NudJ interacts with single nucleotides, catalysing the dephosphorylation of deoxy and ribo nucleotides in both the di and tri-phosphate forms (Xu et al. 2006). This contrast suggests that whilst the antimutator phenotypes of these strains are similar, the molecular mechanisms responsible are distinct.

A more similar case to *nudJ* is *purB* in *E. coli*, where a temperature sensitive mutation (*purB^ts^*) provides an antimutator phenotype (Geiger and Speyer 1977; though see Schaaper and Dunn 2001). Like NudJ, PurB is involved in nucleotide metabolism; PurB plays an essential role in purine biosynthesis (Zhang et al. 2008). Interestingly, *purB* and *nudJ* are only ∼1,800 bp apart in the *E. coli* genome and are also commonly co-expressed (Szklarczyk et al. 2022). This raises the possibility that the antimutator phenotypes of Δ*nudJ* and *purB^ts^* may share a common cause.

The mutation rate reduction seen in Δ*mfd* strains (2- to 5-fold) is similar to our observations in Δ*nudJ* (2-6 fold). When compared to the >100 fold increases in mutation rate seen in some mutator strains these changes are modest. However, dramatic reductions in the ability of *Salmonella typhimurium* to evolve resistance to multiple antibiotics was seen in the absence of *mfd* (Ragheb et al. 2019); demonstrating that small mutation rate decreases can have a large impact on evolutionary trajectories. In that study, genetic mutator strains were frequently observed when the wildtype strain evolved in the presence of trimethoprim however, the evolution of a mutator phenotype was never observed in the Δ*mfd* antimutator. Therefore, antimutators are not only less likely to access mutations providing antibiotic resistance, but are also less likely to access mutations leading to a mutator phenotype, which can facilitate resistance evolution (Couce et al. 2015; Gifford et al. 2023; Elgrail et al. 2024). The logic behind this finding is that the greater a strain’s mutation rate, the greater its chance of accessing mutation rate modifying alleles. This is reflected by frequent observations of both positive (Gifford et al. 2023; Elgrail et al. 2024) and negative (McDonald et al. 2012; Wielgoss et al. 2012; Sprouffske et al. 2018) mutation rate modifying alleles in mutator strains during experimental evolution.

Mutator strains, with elevated mutation rates, often show dramatic changes in mutational spectra which reflect the role of the disrupted gene. For example, in Δ*mutT* strains, oxidised dGTP is no longer removed from the cytoplasm, leading to A:G mispairing events, which result in many A>C transversion mutations (Maki and Sekiguchi 1992). As well as having evolutionary implications, our finding of a significantly altered mutational spectrum in Δ*nudJ* provides hypotheses as to the mechanism of NudJ’s role in mutagenesis. Our finding that *nudJ* deletion specifically decreases the rate of A>G transitions indicates that the presence of NudJ preferentially enables these mutations. An elevated rate of A>G transitions is also observed in *E. coli* starved of the nucleotide thymine and this has been attributed to imbalances in the dNTP pool (Kunz and Glickman 1985). Consistent with the hypothesis that the mutagenic effects of NudJ act via imbalances of the nucleotide pool, we find significantly reduced concentrations of ATP in the *nudJ* deletant strain (Figure S4).

It is possible the the antimutator phenotype of Δ*nudJ* arrises through the same mechanism as the antimutagenic effect of guanosine supplementation reported by (Novick and Szilard 1952); which may or may not be associated with the mutational effects of thymine starvation (Kunz and Glickman 1985) and the ‘dNTP mutator’ effect (Gon et al. 2011). Among 17 deoxy- and ribonucleosides tested by Xu et al. (2006), NudJ shows greatest hydrolase activity towards GTP, it is therefore plausible that in the absence of NudJ more guanosine will accumulate in the cell. Gudkov et al. (2006) show that, in the presence of *γ*-irradiation or heating, guanosine can prevent the formation of 8-oxoguanine (8-OG) and the deamination of cytosine in DNA. These results suggest that guanosine will reduce the rates of multiple mutational classes, consistent with our observation that all mutational classes are suppressed, to differing extents, in Δ*nudJ* (Figure S9). We suggest that future work should measure individual nucleotide, dNDP, dNTP and NTP molecule concentrations in Δ*nudJ* and the wildtype to specifically quantify any imbalance in these nucleotide pools potentially using HPLC-based methods as in Buckstein et al. (2008) & Varik et al. (2017). This will allow us to develop more specific hypotheses as to the mechanism(s) by which NudJ impacts mutagenesis.

Aside from its nucleoside dephosphorylase activity, NudJ (along with Nud B, F & I) is also known to dephosphorylate DMAPP to DMAP which is used to synthesise prenylated flavin mononucleotide (prFMN) (Wang et al. 2018). prFMN is required by UbiD-like enzymes for the biosynthesis of ubiquinone (Wang et al. 2018). This link between antimutators Δ*nudJ,* Δ*nudB* and Δ*nudF* and the known antioxidant ubiquinone (Søballe and Poole 2000) is intriguing given the known role of oxidative stress in determining mutation rates (e.g. (Lagage et al. 2022; Carvajal-Garcia et al. 2023; Green et al. 2024)). NudJ, NudF and NudB are also known to hydrolyse isopentenyl diphosphate (IPP) to 3-methyl-3-butenol (Chou and Keasling 2012), (Figure S2). IPP and DMAPP (both degraded by NudJ) are combined to form geranyl-PP by IspA (Fujisaki et al. 1986), (Figure S2). This may explain the phenotypic reversion associated with a secondary *ispA* mutation in a Δ*nudJ* strain, while *nudJ* deletion may increase intracellular geranyl-PP concentrations, the disruption of *ispA* will reduce the genranyl-PP pool, countering the effect of *nudJ* deletion. However, as *ispA* and *csgF* are disrupted in the same Δ*nudJ* strain it is possible that either one of these secondary mutations, or the combined effects of both, are responsible for the mutation rate reversion (Figure 2, Figure S5).

The contributions of WaaZ and potentially of CsgF to mutagenesis in the Δ*nudJ* background also remains to be explored. Both are disrupted by IS insertions in our revertant Δ*nudJ* strains, which may have an impact on not only their own transcription but that of the other genes in their operons (Jubelin et al. 2005; Lee et al. 2009; Kanai et al. 2022). We suggest that the reversal of the Δ*nudJ* phenotype by these secondary mutations in the outer membrane and extracellular matrix may operate by altering permeability of the cell to nucleotides, reversing the effect of the *nudJ* deletion on the internal nucleotide pool (Goyal et al. 2014; Wang et al. 2021). In line with this, we find that while the internal ATP concentration is decreased in Δ*nudJ*, these revertant strains show a return to wildtype or greater internal ATP concentrations (Figure S4).

As well as identifying novel *E. coli* antimutators, this study also highlights the general importance of considering not only mutation rates but also mutational spectra in predicting evolutionary outcomes. Here we use rifampicin resistance mutations to explore the effect of mutational spectra on fitness outcomes. The highly constrained nature of rifampicin resistance evolution is reflective of many antibiotics for which resistance can be gained by target site modification. Because of the more modest mutation rate changes seen in antimutators, as opposed to mutators, the effect of mutational spectrum is likely to be more pronounced. This makes the potential for rational manipulation of mutational spectrum by antievolution drugs, designed to reduce mutation rates, a feasible method for increasing their success. We therefore suggest that future work should assess the mutational spectra within known antimutators to build a more general picture of antimutator phenotypes. The environmental responsiveness of both mutation rates and spectra demonstrated in this study will be a hurdle to the development of such drugs. It is therefore important for future research on mutation rate manipulation to consider both environmental plasticity as well as the relationship between mutational spectrum and fitness outcomes in the environment of interest (e.g. an infection under antibiotic treatment). By bringing together all of these facets shaping evolvability, we will improve our ability to predict evolutionary outcomes, opening up the possibility of using this knowledge to manipulate the course of evolution.

## Materials & Methods

### Strains and Media

The wildtype BW25113 is the ancestor of Δ*waaZ,* Δ*thiM* and Δ*nudBCDEFGHIJK* which were sourced from the Keio collection (Baba et al. 2006). All knockouts in the Keio collection appear twice; Δ*waaZ* was sourced from the first available knockout, Δ*thiM* from the second available knockout and all stocks of Δ*nudBCDEFGHIJK* from the first available knockout aside from Δ*nudJ* + fs *ispA* + IS5 *csgF* which derives from the second *nudJ* knockout. Wildtype MG1655 was kindly provided by Karina Xavier and is the ancestor to all *rpoB* mutants used for growth assays. These MG1655 *rpoB* mutants were collected and sequenced by Gifford et al. (2024).

We used Milli-Q water for all media, all chemicals are supplied by Sigma-Aldrich unless stated otherwise. LB medium contained: 10 g of NaCl (Thermo Fisher Scientific), 5 g of yeast extract (Thermo Fisher Scientific) and 10 g of tryptone (Thermo Fisher Scientific) per litre. DM medium contained 0.5 g of C_6_H_5_Na_3_O_7_ ·2H_2_O, 1 g of (NH_4_)2SO_4_ (Thermo Fisher Scientific), 2 g of H_2_KO_4_P and 7 g of HK_2_O_4_P· 3H_2_O per litre; 100 mg L^-1^ MgSO_4_ ·7H_2_O (406 µmol), 4.4 μg L^-1^ thiamine hydrochloride and glucose to the desired concentration (80-1000 mg) were added to DM after autoclaving. M9 medium contained 6.8 g of Na2HPO4, 3 g of KH2PO4, 0.5 g of NaCl and 1 g of NH4Cl per liter; 100uM CaCl2.2H2O, 2mM MgSO4.7H2O, 3µM thiamine hydrochloride, 11.2mM glucose and 0.2% casamino acids were added after autoclaving. Selective tetrazolium arabinose agar (TA) medium contained 10 g of tryptone, 1 g of yeast extract, 3.75 g of NaCl and 15 g bacto agar per litre; after autoclaving 3 g of arabinose and 0.05 g of 2,3,5-triphenyl-tetrazolium chloride were added per litre, this was supplemented with freshly prepared rifampicin (50 µg ml^−1^) dissolved in 1mL of methanol. For all cell dilutions sterile saline (8.5 g L^-1^ NaCl) was used.

### Fluctuation assays

Fluctuation assays were conducted as described in (Krašovec et al. 2019). Briefly; an ice scrape from glycerol stock was incubated in LB for 4-7 hrs (until visibly turbid). A 1000 fold dilution then used for transfer to 10mL overnight cultures in DM with the appropriate glucose concentration for acclimatisation (80-1000 mg.L^-1^). The density of these overnight cultures was then measured by OD and all cultures were diluted to a hypothetical OD of 3×10^-6^, equating to ∼3000 CFU per mL, in fresh media. Large initial cultures were split into 19-20 parallel cultures of 0.5-1.25mL in deep 96 well plates and incubated for 24 hours. For each treatment 3 ‘Nt’ wells were taken and diluted appropriately before plating on non-selective TA agar to determine the final population size and density reached. The remaining wells were plated in their entirety on selective TA agar plates containing 50mg.L^-1^ rifampicin and dried in a sterile biosafety cabinet. ‘Nt’ plates were incubated for 24 hours and selective plates for 44-48 hours before colonies were counted. Estimates of mutational events were calculated using the maximum likelihood method as implemented by R package flan (Mazoyer et al. 2021). Because of the known effect of density on microbial mutation rates we estimate mutation rates for all strains at the mean density of the dataset (Figure 1, Figure S1) (Krašovec et al. 2014; Krašovec et al. 2017; Krašovec et al. 2018; Green et al. 2024).

Intracellular ATP was assayed at the end of most fluctuation assays (308 / 402) via luminescence using a Promega GloMax luminometer and the Promega Bac-Titer Glo kit, according to the manufacturer’s instructions. We measured the luminescence of each culture 0.5 and 510 s after adding the Bac-Titer Glo reagent and calculated net luminescence as luminescence_510s_ − luminescence_0.5s_.

Liquid based fluctuation assays were used in comparing the mutation rates of Δ*waaZ* and the wildtype (Figure S6). The protocol remained the same as above aside from the identification of mutants in selective wells. Rather than plating the wells on rifampicin agar, fresh LB was added to the wells at this point along with rifampicin to a final concentration of (50mg.L^-1^). After 2 days of further incubation, OD measurements were taken for all selective wells, OD>0.2 was categorized as presence of a mutant and OD≤0.2 as absence. Mutation rates were estimated using the P0 method as implemented by R package flan (Mazoyer et al. 2021).

### PCR and Sanger Sequencing

rifR mutants were picked from selective plates at the end of low density fluctuation assays (80mg.L^-1^ glucose) with the BW25113 wildtype, Δ*nudJ* or Δ*nudJ* + IS1 *waaZ* and grown to turbidity (4-5 hrs) in LB before freezing in 18% glycerol at -80℃. Mutants were then streaked from glycerol stocks onto TA agar and one colony was picked into 25µL molecular grade H_2_O and mixed to break the cells by osmotic lysis, this solution was used as the DNA template. 25µL PCR reactions contained 2.5µL DNA template, 0.125µL Phusion High-Fidelity DNA Polymerase (M0530S), 0.25µL 500 μM forward and reverse primers, 1.25µL DMSO, 5µL buffer, 0.5µL 10mM dNTPs and 15.125µL nuclease free H_2_O. Forward primer 5′-ATGATATCGACCACCTCGG-3′ and reverse primer 3′-TTCACCCGGATACATCTCG-5′ were used for all reactions as in (Gifford et al. 2024). PCR was run with the following protocol: (i) initial denaturation (98°C for 5 min); (ii) denaturation (98°C for 10 s); (iii) annealing (55°C for 30 s); (iv) extension (72°C for 1 min); (v) repeat steps 2 to 4 for 30 cycles; (vi) final extension (72°C for 5 min); and (vii) hold at 4°C. PCR product verification was performed by gel electrophoresis carried out on 1% agarose TAE gel with 0.1% SybrSafe stain. PCR products were submitted to Source BioScience for PCR product clean-up and sequencing (Source BioScience, Cheshire). Each product was submitted with either forward or reverse primer as the sequencing primer.

Raw Sanger sequencing results were converted to trimmed fasta files using R package SangeranayseR (Chao et al. 2021), mutations were then identified by alignment to the reference sequence (EG10894). All mutations were verified by visual inspection of the relevant chromatogram.

Of 464 sequenced isolates 2 had 2 mutations in the RRDR, these 4 mutations were excluded (Strain 1: wildtype position 1604 C>T + 1715 T>A, Strain 2: Δ*nudJ* 1598 T>G + 1604 C>T). This is because we cannot be sure of the order in which these mutations occurred; once a first mutation is acquired in *rpoB* the mutational spectrum for the second mutation may be altered due to pleiotropic effects of *rpoB* mutations (Bergval et al. 2007; Jago et al. 2024). Where 2 peaks were present at a given position we required the primary peak to be over 3x the height of a secondary peak in order to accept the mutation as conclusive, 2 mutations were excluded by this requirement (1 wildtype 1687 A>C, 1 nudJ +waaZ 1687 A>C). If we include the 6 filtered mutations our finding of a significantly altered mutational spectrum in Δ*nudJ* vs wildtype is retained (LR = 17.8, *P* = 0.0262), as are the respective increase and decrease of A>C and A>G mutations in Δ*nudJ* (*χ*^2^ = 10.8 & 7.34, 0.00791 & 0.027 respectively). Our finding of reduced fitness in rifR mutants from the Δ*nudJ* background across all 12 tested environments is also retained (t_DF=1555_ = -4.15, *P* = 3.45^{-5}). 65 of 464 sequenced mutants have no mutation in the RRDR (14% of filtered reads), this is to be expected as rifampicin resistance can be achieved by mutations outside the RRDR (Garibyan 2003). For the wildtype and Δ*nudJ* + IS1 *waaZ* ∼20% of sequenced strains show no mutation in the RRDR (24/116 & 24/126 respectively) whereas for Δ*nudJ* <8% of sequenced strains (17/218) show no mutation in the RRDR. This is consistent with the findings of (Garibyan 2003) who find that, in the wildtype background, 23/24 identified mutations outside the RRDR are A>T or A>G mutations, both proportionally underrepresented in Δ*nudJ* (Figure 3).

The inclusion of block effects significantly improves the fit of a multinomial model of the effect of genotype on mutational spectrum (LR = 33.7, df = 21, *P =* 0.0386). These block effects are likely the result of slight differences in conditions between experimental blocks; for example changes in media composition and growth temperature are known to impact mutational spectrum (Chu et al. 2018; Liu and Zhang 2019). We therefore use a model accounting for block effects when analysing mutational spectrum in order to account for this potentially confounding effect.

### Growth Curves

Growth curves of rifR mutants in the MG1655 background were initiated by incubating 1uL from frozen glycerol stocks in 200µL LB at 37℃ with 200rpm shaking overnight. These cultures were then diluted 200 fold into appropriate fresh media (M9 or LB), with or without rifampicin (50mg.L^-1^) and grown at 25, 37 or 42℃ (giving a total of 12 possible environments). Clear 96-well plates were incubated for 24 hours with OD readings taken every 30 minutes using BMG Omega plate reader. 34 mutants were tested in total (27 of which bear mutations identified in the BW25113 wildtype, Δ*nudJ* or Δ*nudJ* + IS1 *waaZ* strains) with 6 repeats per mutant in each environment. Each mutant appeared twice within each 96 well plate, in order to minimise the effect of greater evaporation in the edge wells on our results no mutant was placed in an edge well twice within a single plate. Summary statistics were computed for each growth curve individually using R package growthcurver (Sprouffske 2020) and means taken across each environment x mutant combination were paired to mutants identified in the BW25113 wildtype, Δ*nudJ* and Δ*nudJ* + IS1 *waaZ* strains for further analysis.

Growth curves of the BW25113 wildtype, Δ*nudJ* and Δ*nudJ* + IS1 *waaZ* were prepared as for a fluctuation assay and grown in minimal DM with either 80 or 1,000 mg of glucose per L. Parallel cultures of 150µL were grown in a shallow 400 µL well plates, and OD600 readings were taken every 10 minutes with shaking at 200rpm for 1 minute prior to each reading.

### Estimation of ΔΔG caused by *rpoB* mutations

ΔΔG of each mutant identified by RRDR sequencing was estimated using FoldX 5.0 (Schymkowitz et al. 2005). Reference protein structure for the RpoB protein was downloaded from alphafold (AF-P0A8V2-F1-v4) (Jumper et al. 2021; Varadi et al. 2023), based on UniProt entry P0A8V2 (Ovchinnikov et al. 1981; Bateman et al. 2022). Reference protein structure for the RNAP holoenzyme complex was downloaded from the Protein Data Bank (Braffman et al. 2019). Before introducing AA substitutions, the RpoB and RNAP protein structures were repaired using the FoldX command ‘RepairPDB’ with options: *--ionStrength=0.05 --pH=7 -- water=CRYSTAL --vdwDesign=2 --pdbHydrogens=FALSE.* FoldX command BuildModel was then used with default settings to estimate the ΔΔG resulting from each observed AA substitution for both RpoB and RNAP separately.

### Whole Genome Sequencing

Knockouts Δ*nudB,* Δ*nudF,* Δ*nudI* and the wildtype BW25113 were sequenced to ≥30x coverage with illumina short reads by MicrobesNG (https://microbesng.com); verifcation that strains carried the correct gene knockouts and no additional genic mutations was performed using breseq version 0.38.1 with default parameters. The wildtype BW25113 genome was used as a reference (Grenier et al. 2014) as modified by (Gifford et al. 2023) to include improved IS annotation.

All Δ*nudJ* strains were sequenced by MicrobesNG (https://microbesng.com) using their hybrid service combining ≥30x coverage with illumina short reads and ≥50x coverage with Oxford nanopore technology long reads. All mutations were predicted using breseq (Deatherage and Barrick 2014) with the wildtype BW25113 genome as a reference (Grenier et al. 2014) modified to include improved IS annotation by (Gifford et al. 2023). Mutations were also verified by using short reads from one Δ*nudJ* strain as a query and the full assembled genome from another Δ*nudJ* strain as the reference, as provided by MicrobesNG (https://microbesng.com). Both methods gave identical predictions.

### Bootstrapping fitness differences between antibiotic resistance mutations accessed by Δ*nudJ* vs the wildtype

We analyzed the impact of mutational spectra on antibiotic resistance costs for nalidixic acid (Nal) and trimethoprim (Tmp) resistant mutants Figure S16. Strains one base-pair substitution removed from the wildtype from Harmand et al. (2017) (Nal) and Palmer et al. (2015) (Tmp) were included. Separately for each antibiotic 100 mutants were sampled, with replacement, following the mutational spectra observed in Δ*nudJ* and the wildtype among rifampicin resistant mutants in this study Figure 3. The sampling probability for each mutation type was proportional to its frequency in the rifampicin resistance spectrum. For example, in the case of the wildtype, 28% of rifR mutants carry A>C transversions while only 10% are C>T transitions therefore 2.8 A>C mutants will be sampled for each C>T mutant sampled. The mean selection coefficient (nalidixic acid) or AUC of a 30hr growth curve was then calculated for the set of 100 mutants sampled by Δ*nudJ* and the wildtype in each tested environment and the difference between these mean values recorded. This bootstrapping procedure was repeated 1000 times for each antibiotic. For nalidixic acid 13 unique mutations each causing a distinct amino acid substitution were included while for trimethoprim 8 unique mutations, one in the promoter region and seven in the coding region (two of which result in the same AA substitution), were included.

This approach does not consider either the presence of mutational hotspots in any of the target genes considered (*rpoB, gyrA* & *folA* for rif, nal & tmp respectively). It also does not account for differences in the spectrum of available mutations to each antibiotic (for example 50% of possible resistance mutations to one drug may be A>G transitions while this type of mutation may account for only 10% of possible resistance mutations to another drug). These limitations are, to some extent, a result of incomplete knowledge of the entire set of mutational targets for any of these drugs. Even with extensive mutant selection and sequencing, mutational ‘coldspots’ may remain unobserved (Sun and Lind 2023). Despite these limitations, this method provides a valuable first approximation of how the costs of antibiotic resistance may be influenced by the specific mutational spectra of Δ*nudJ* vs the wildtype.

### Statistical Methods

All statistical models associated with this study are detailed in the Supplementary Statistical Methods document.

## Supporting information

Supplementary Statistics

## Abbreviations

IS: Insertion sequence
SNP: Single nucleotide polymorphism
DMAPP: Dimethylallyl-diphosphate
DMAP: Dimethylallyl-monophosphate
prFMN: Prenylated flavin mononucletide
TPP: Thiamin phyrophospate
HMP-PP: 4-amino-2-methyl-5-hydroxymethylpyrimidine pyrophosphate
HET: 4-methyl-5-hydroxyethylthiazole
DAMP: Density associated mutation rate plasticity
rifR: rifampicin resistance
RRDR: Rifampicin resistance determining region of *rpoB*
AA: Amino acid
DFE: Distribution of fitness effects
AUC: Area under the curve
ΔΔG: Change in the Gibbs free energy of folding
RNAP: RNA polymerase
CFU: Colony forming units
PCR: Polymerase chain reaction

## Acknowledgements

Thanks to Sandy McLennan for helpful correspondence concerning the biochemistry of the Nudix hydrolase enzymes, to Karina B. Xavier for kindly sharing the MG1655 wildtype strain and to MicrobesNG (http://www.microbesng.com) for their genome sequencing services.

## Data Availability

Data and analysis code are archived with the Open Science Framework at osf.io/mu2ek.

