## Supplementary Figures & Tables for "Evolutionary potential of the *Escherichia coli* antimutator Δ*nudJ* is reduced via altered mutational spectrum"

### Supplementary Materials

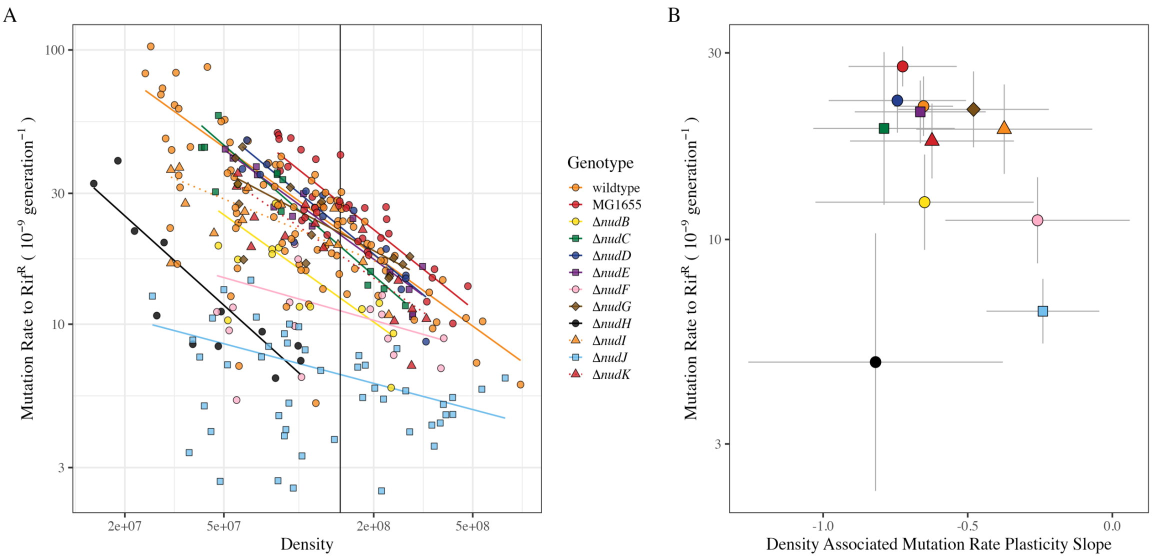

Figure S1: **Data underlying mutation rate estimates for Nudix hydrolase knockouts.** A) Mutation rate to rifampicin resistance (mutational events per division (x$10^9$)) plotted as a function of final population density (CFU per mL). Raw data points are normalised by subtracting random effects associated with experimental block (see supplementary Statistical Methods document). B) Coefficients with 95% CI from a regression model fitting mutation rate as a function of genotype and population density with interaction (Methods). Wildtype BW25113 is shown as orange circles ($N_{\text{Fluctuation Assays}}$ = 92, $N_{\text{Parallel Cultures}}$ = 1613), wildtype MG1655 is shown as red circles ($N_{FA}$ = 35, $N_{PC}$ = 567), **Δ***nudB* is shown as yellow circles ($N_{FA}$ = 14, $N_{PC}$ = 228), **Δ***nudC* is shown as green squares ($N_{FA}$ = 12, $N_{PC}$ = 193), **Δ***nudD* is shown as dark blue circles ($N_{FA}$ = 14, $N_{PC}$ = 227), **Δ***nudE* is shown as purple squares ($N_{FA}$ = 14, $N_{PC}$ = 228), **Δ***nudF* is shown as pink circles ($N_{FA}$ = 16, $N_{PC}$ = 259), **Δ***nudG* is shown as brown diamonds ($N_{FA}$ = 14, $N_{PC}$ = 226), **Δ***nudH* is shown as black circles ($N_{FA}$ = 12, $N_{PC}$ = 195), **Δ***nudI* is shown as orange triangles ($N_{FA}$ = 14, $N_{PC}$ = 225), **Δ***nudJ* is shown as light blue squares ($N_{FA}$ = 47, $N_{PC}$ = 1089), **Δ***nudK* is shown as red triangles ($N_{FA}$ = 14, $N_{PC}$ = 226).

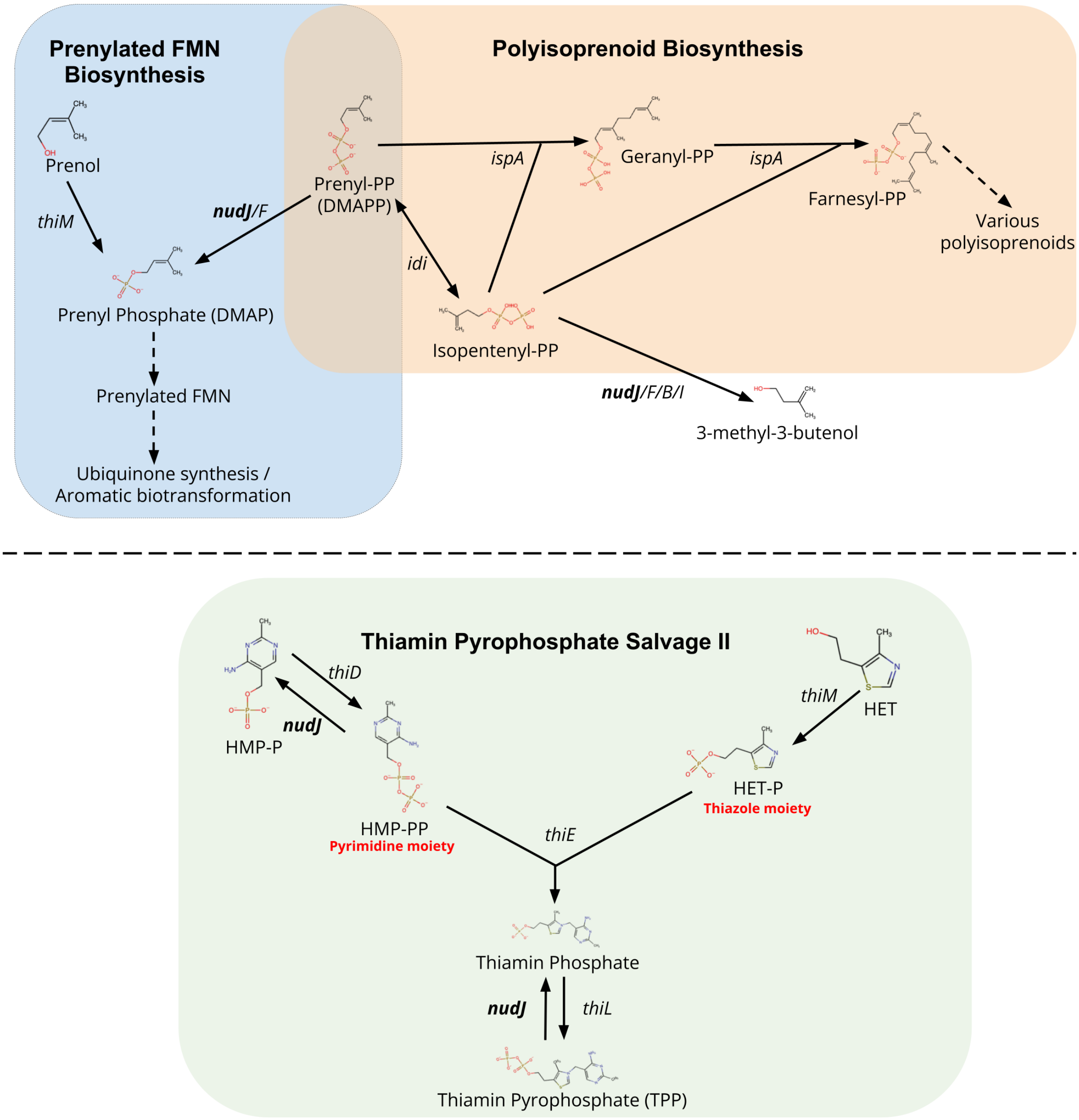

Figure S2: **Simplified diagram of the comparative roles of ThiM and NudJ.** In the prenylated FMN biosyhtnesis pathway both NudJ & ThiM produce DMAP, enabling prenylated FMN production. However, in thiamin pyrophosphate salvage ThiM contributes positively to TPP production while NudJ antagonises TPP production. The antagonistic roles of NudJ and IspA in polyisoprenoid biosynthesis are also included. See main text for references associated with all reactions shown. All chemical structures were generated using the RCSB PDB Chemical Sketch Tool (https://www.rcsb.org/chemical-sketch).

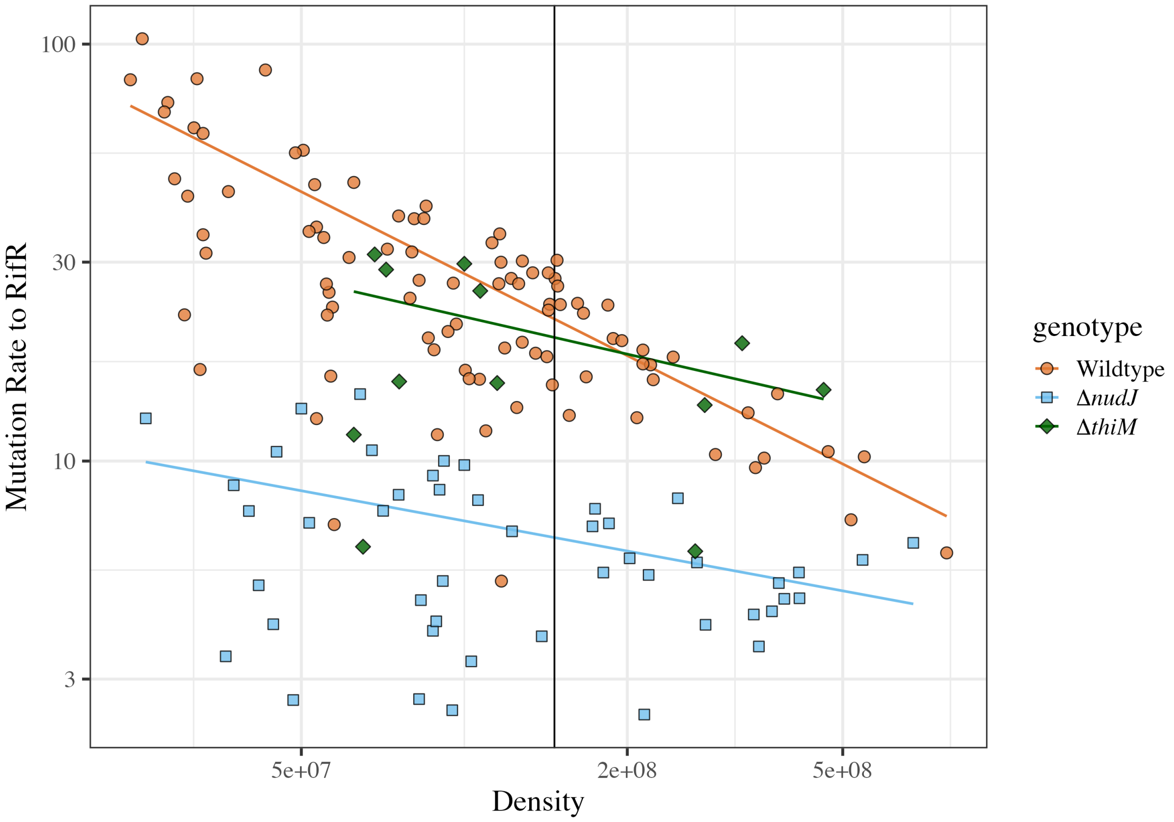

Figure S3: **Mutation rate of *thiM* deletant.** Mutation rate to rifampicin resistance (mutational events per division (x10$^9$)) plotted as a function of final population density (CFU per mL). Wildtype is shown as orange circles ($N_{\text{Fluctuation Assays}}$ = 92, $N_{\text{Parallel Cultures}}$ = 1613, **Δ***nudJ* as blue squares ($N_{FA}$ = 47, $N_{PC}$ = 1089 and **Δ***thiM* as green diamonds ($N_{FA}$ = 12, $N_{PC}$ = 204). Raw data points are normalised by subtracting random effects associated with experimental block (see supplementary Statistical Methods document). Lines of best fit are shown with the relevant colour for each genotype as fitted by Statistical model 1.

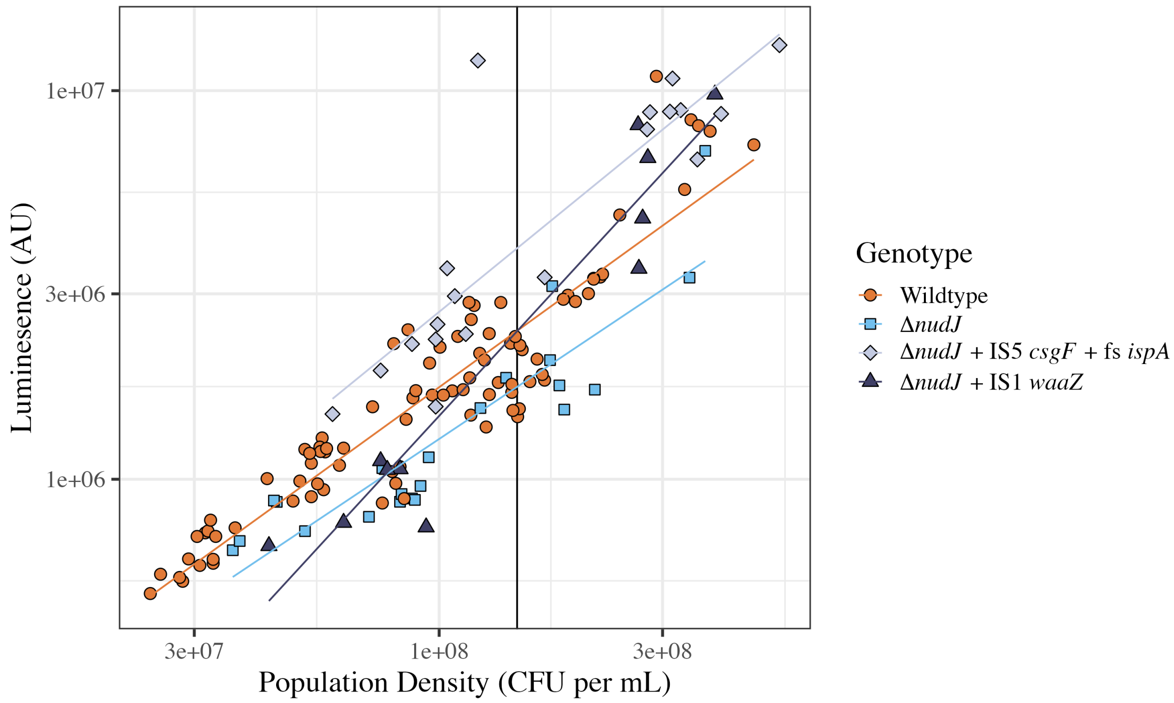

Figure S4: **ATP based luminescence assay.** Luminescence in arbitrary units (AU) is plotted as a function of population density. All luminescence measurements are normalised by subtracting random effects of experimental block and plate as well as fixed effects of experimenter identity, as estimated by Statistical Model 2. Lines of best fit are shown as estimated by Statistical Model 2. The wildtype is shown as orange circles ($N = 88$), **Δ***nudJ* is shown as pale blue squares ($N = 22$), **Δ***nudJ* + IS5 *csgF* + fs *ispA* is shown as grey diamonds ($N = 19$) and **Δ***nudJ* + IS1 *waaZ* is shown as dark blue triangles ($N = 11$). Note log-log axes.

A

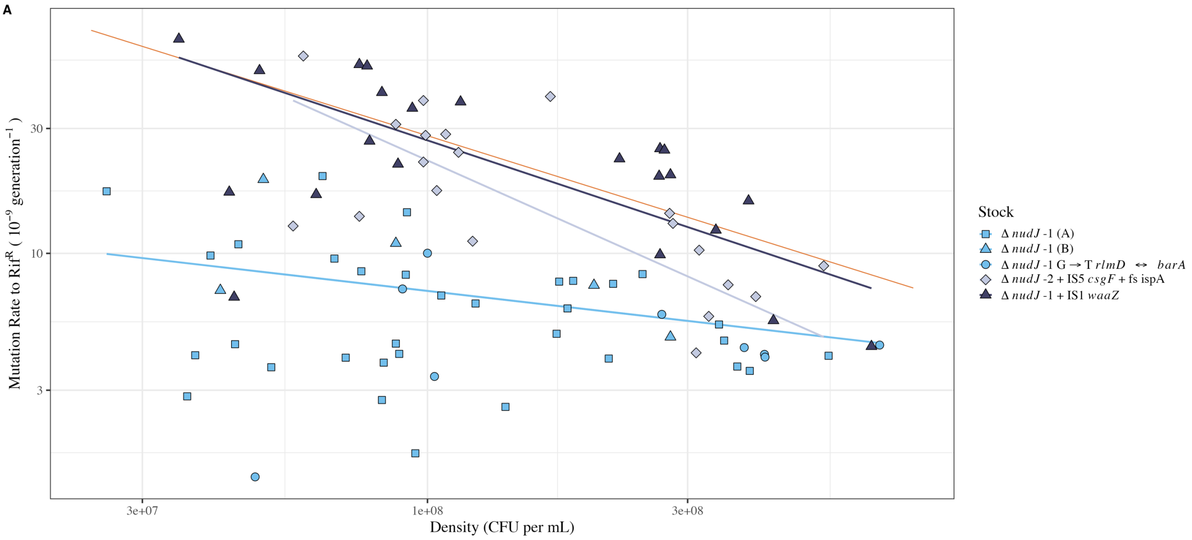

B

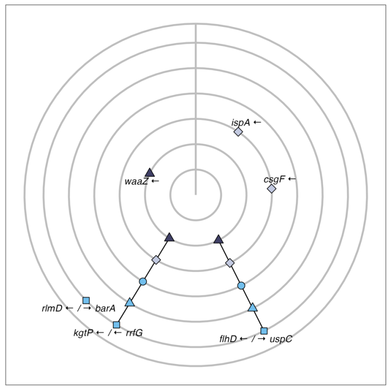

C

| Strain | position | mutation | annotation | gene |
|---|---|---|---|---|
| Δ *nudJ* -2 + IS5 *csgF* + fs ispA | 436547 | +CG | coding (11/900 nt) | ispA ← |
| Δ *nudJ* -2 + IS5 *csgF* + fs ispA | 1097379 | IS5 (–) +4 bp | coding (202-205/417 nt) | csgF ← |
| Δ *nudJ* -1 G → T *rlmD* ← / → *barA* | 2908408 | G→T | intergenic (-49/-8) | rlmD ← / → barA |
| Δ *nudJ* -1 + IS1 *waaZ* | 3793366 | IS1 (–) +9 bp | coding (183-191/852 nt) | waaZ ← |
| All | 1972967 | IS5 (+) +4 bp | intergenic (-513/-264) | flhD ← / → uspC |
| All | 2719426 | T→C | intergenic (-321/+2) | kgtP ← / ← rrfG |

Figure S5: **Mutation rates of 5 Δ*nudJ* freezer stocks.** **Δ***nudJ*-1 and **Δ***nudJ*-2 denote the 2 independently created *nudJ* deletant strains in the Keio collection. **Δ***nudJ*-1 (A) & (B) indicate the 2 separate freezer stocks taken from Keio deletant **Δ***nudJ*-1 with no secondary mutations. All other stocks are identified by their secondary mutations. A) Mutation rate to rifampicin resistance (mutational events per division (x$10^9$)) plotted as a function of final population density (CFU per mL). The orange line shows fitted values for the wildtype as a reference. 2 stocks show no off target mutations and are shown as light blue squares (**Δ***nudJ*-1 (A), $N_{\text{Fluctuation Assays}} = 19$, $N_{\text{Parallel Cultures}} = 306$) and light blue triangles (**Δ***nudJ*-1 (B), $N_{FA} = 0$, $N_{PC} = 0$), one stock had no off target genic mutations and an intergenic G>T transversion between *rlmD* and *barA* is shown as light blue circles (**Δ***nudJ*-1 + G⟶T *rlmD* ⟵/⟶ *barA*, $N_{FA} = 0$, $N_{PC} = 0$), the two stocks with off target genic mutations are shown as grey diamonds (**Δ***nudJ*-2 + *ispA* fs + *csgF* IS5, $N_{FA} = 0$, $N_{PC} = 0$) and dark blue triangles (**Δ***nudJ*-1 + *waaZ* IS1, $N_{FA} = 0$, $N_{PC} = 0$). Levels are combined for the 3 stocks with no off target genic mutations, improving the fit of the model (Statistical Model 1). Note log-log axis scales in panel A. B) Visualisation of identified mutations in 5 **Δ***nudJ* stocks. Vertical line represents the origin of replication. C) Table of secondary mutations identified in 5 **Δ***nudJ* stocks. Mutations are identified with reference to the wildtype BW25113 and positions are given for the reference genome. All predictions made with breseq (Deatherage and Barrick 2014).

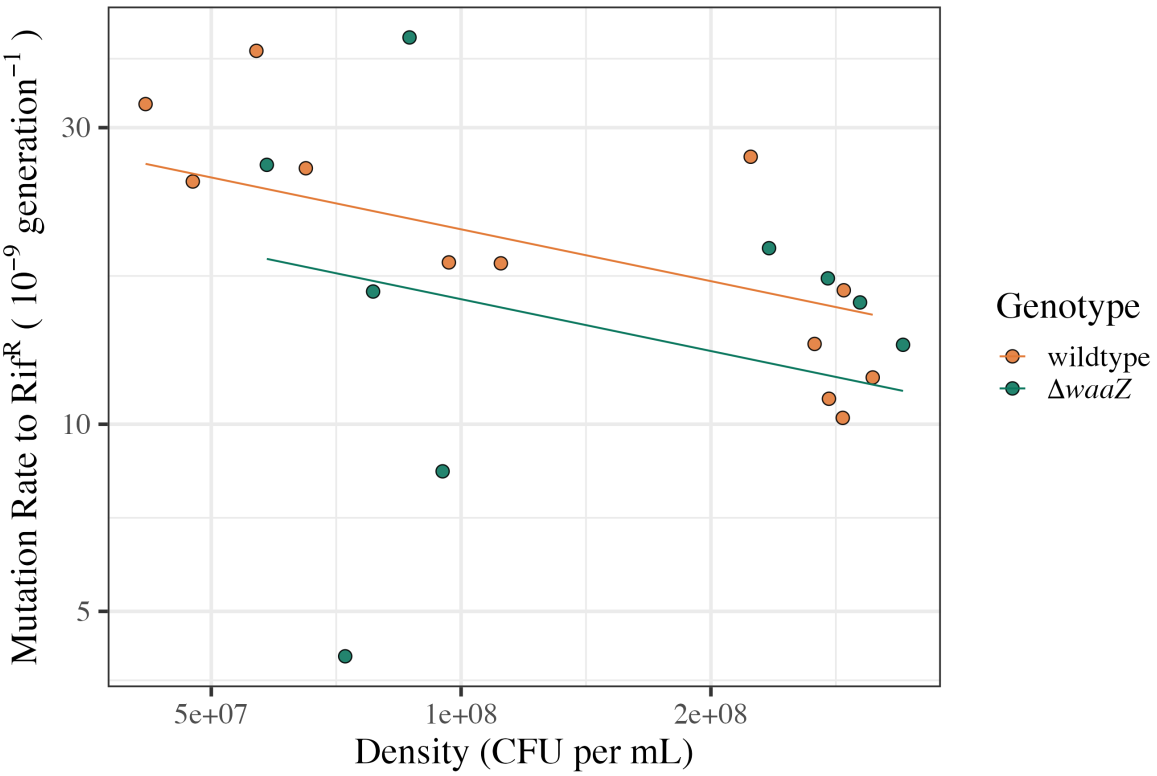

Figure S6: **Mutation rate of Δ*waaZ* is not elevated compared to wildtype BW25113** Mutation rate to rifampicin resistance (mutational events per division (x$10^9$)) plotted as a function of final population density (CFU per mL). Genotype is indicated by colour: orange circles = wildtype, *N*=12; green circles = **Δ***waaZ*, *N*=9. Mutation rate is estimated using a liquid based fluctuation assay to rifampicin resistance (Methods). Note log-log axes.

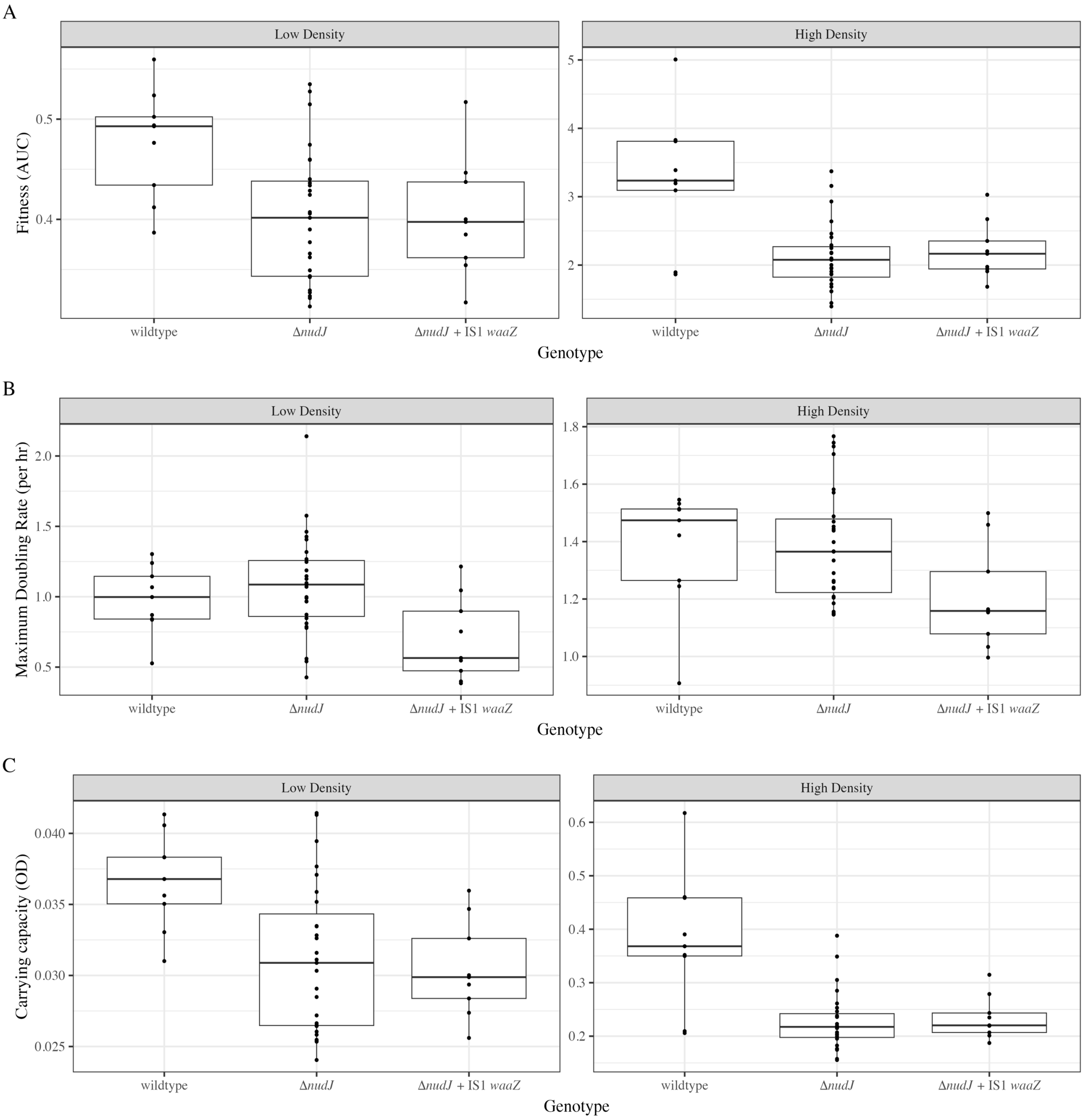

Figure S7: **Loss of fitness in Δ*nudJ* strains results from lower carrying capacity** Data from growth curves with 3 biological replicates each with 3 technical replicates for each genotype aside from **Δ***nudJ* for which 9 biological replicates each with 3 technical replicates were collected.

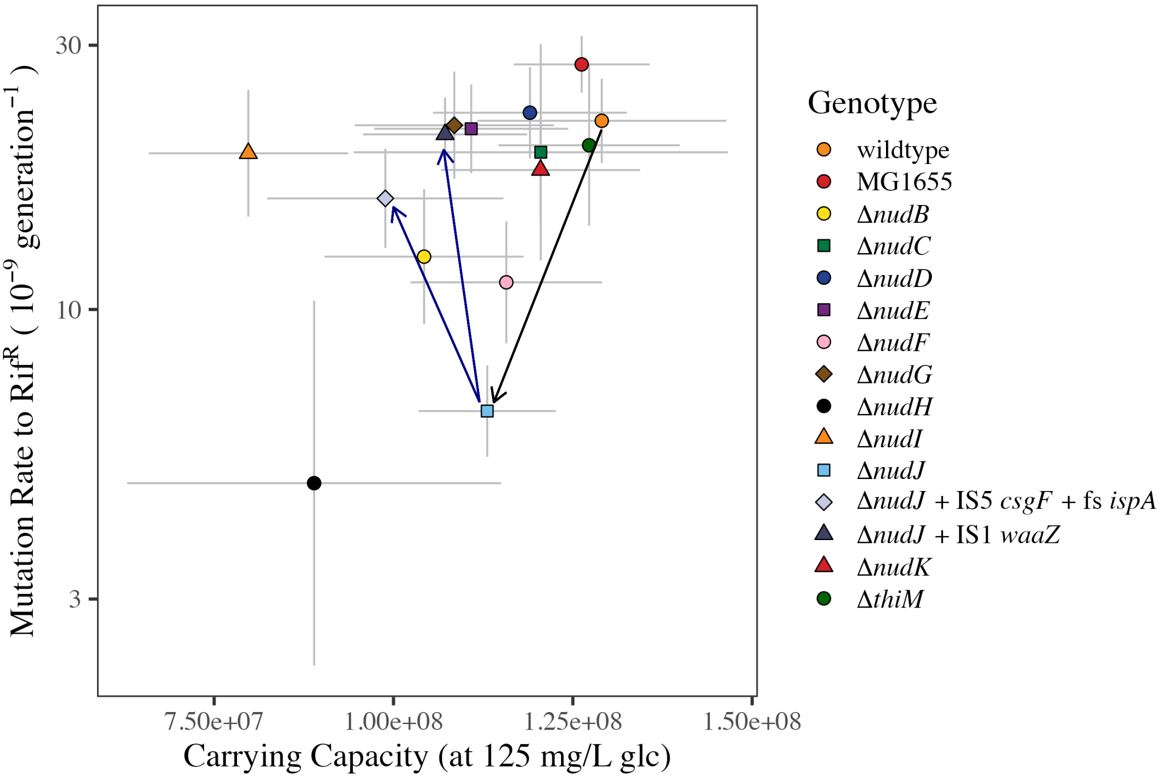

Figure S8: **Mutation rate against carrying capacity: any growth defect in Δ*nudJ* is not recovered by secondary mutations in *waaZ*/*csgF*+*ispA*.** Carrying capacity at 125mg per L glucose is estimated from final CFU counts on non-selective agar from 24 hour growth. Black line shows the reduction in carrying capacity and mutation rate caused by *nudJ* deletion. Blue arrows show that mutation rate, but not carrying capacity, reverts to the wildtype phenotype after secondary mutations. Carrying capacity at 125mg/L glucose fitted by Statistical Model 8 and mutation rate at mean density fitted by Statistical Model 1. Error bars show 95% CI for these parameter estimates.

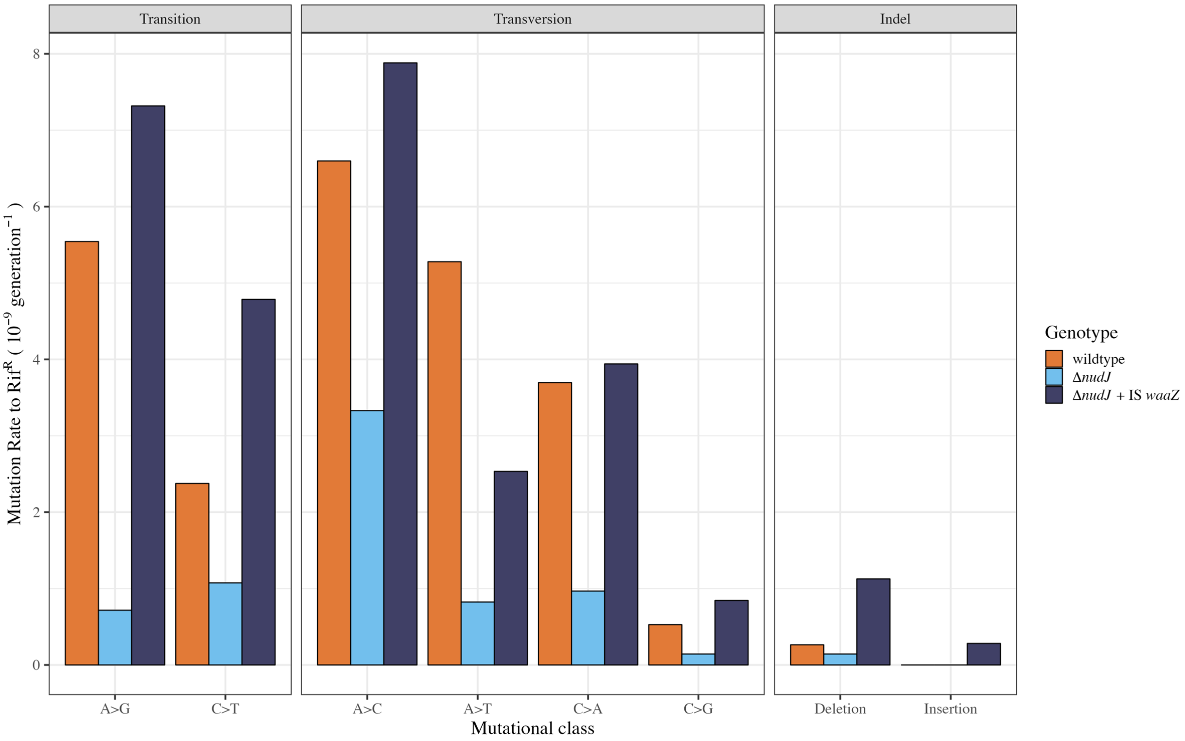

Figure S9: **Mutation rates are reduced by *nudJ* deletion, to a varying extent, across all mutational classes.** For each genotype the mutation rate to rifampicin resistance for a given mutational class is calculated as the proportion of mutants falling into this class (as shown in Figure 3) multiplied by the average mutation rate across the low density fluctuation assays from which sequenced mutants were collected (Mutation rates are: *nudJ* knockout - $7.20 \times 10^{-9}$; wildtype - $2.43 \times 10^{-8}$; *nudJ* knockout + *waaZ* IS1 - $2.87 \times 10^{-8}$). This is a simplification which ignores the rifR mutants carrying no mutation in the RRDR which were identified in all 3 genetic backgrounds (discussed in the methods section).

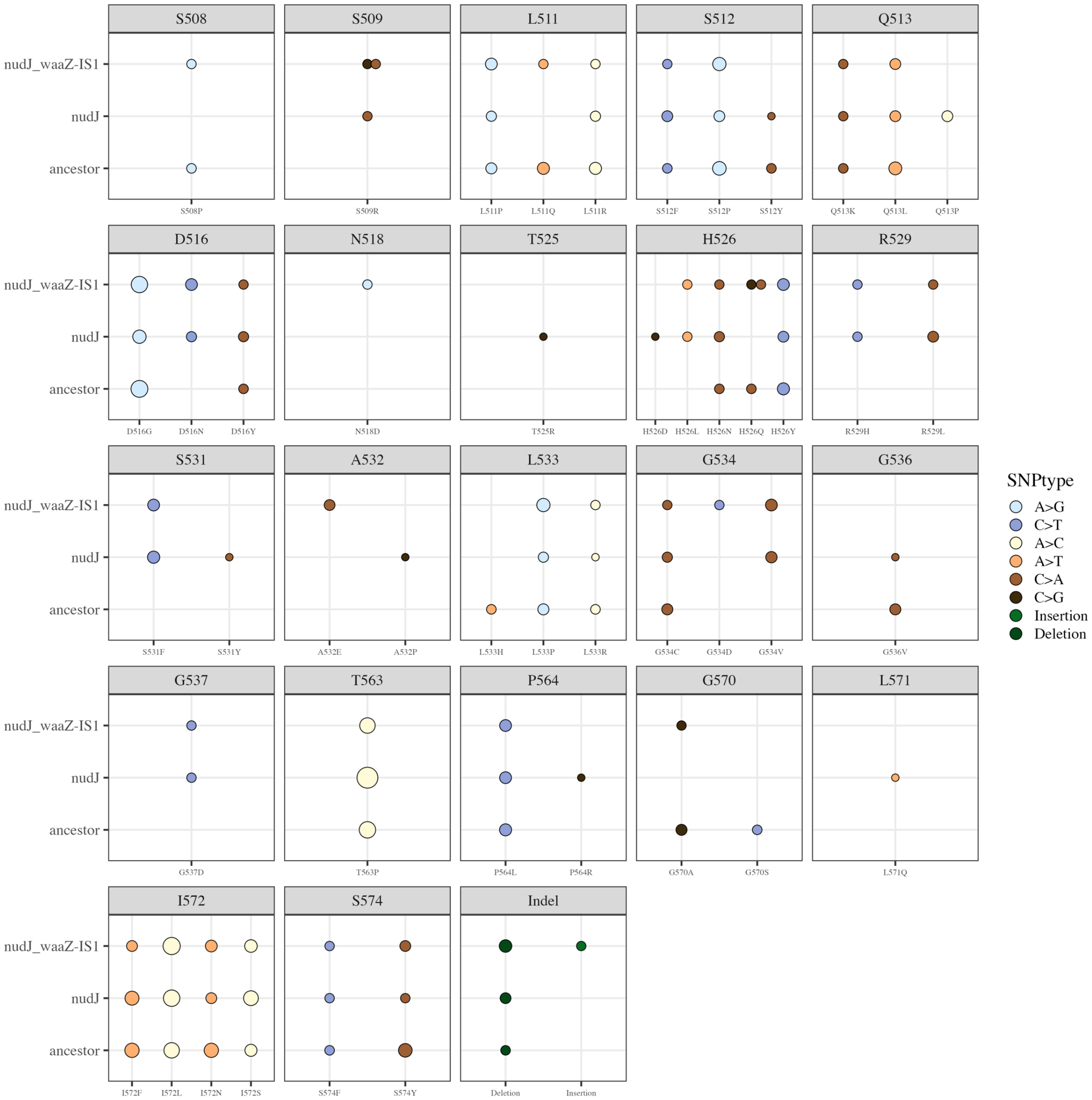

Figure S10: **All observed AA changes in Sanger sequencing of the rifR determining region.** Each subplot indicates a targetted amino acid with the observed replacement amino acid given on the x axis, and the strain in which the given AA substitution was observed given on the y axis. Points are coloured by mutational class and size indicates the proportion of mutations in the given strain accounted for by the given AA substitution (range 0.5% - 24%).

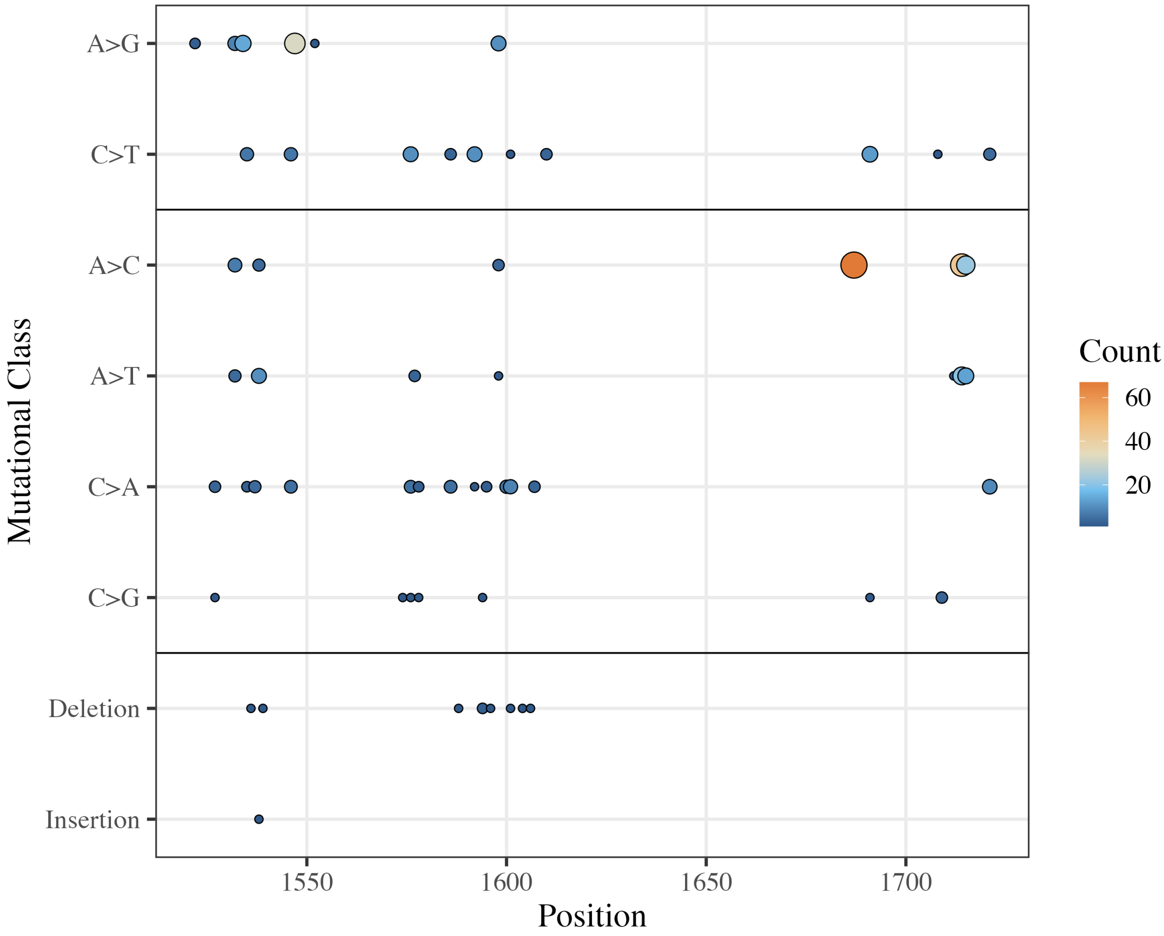

Figure S11: **All observed mutations observed in the wildtype BW25113, Δ*nudJ* and Δ*nudJ* + IS1 *waaZ* backgrounds combined.** Mutations are plotted by position in the genome along the x axis and divided into the 8 observed mutational classes along the y axis. The three sections from top to bottom are transitions, transversions & indels. Each point represents a unique mutation with size and colour indicating the number of times the given mutation was observed among a total of 395 mutations (range 1 - 67).

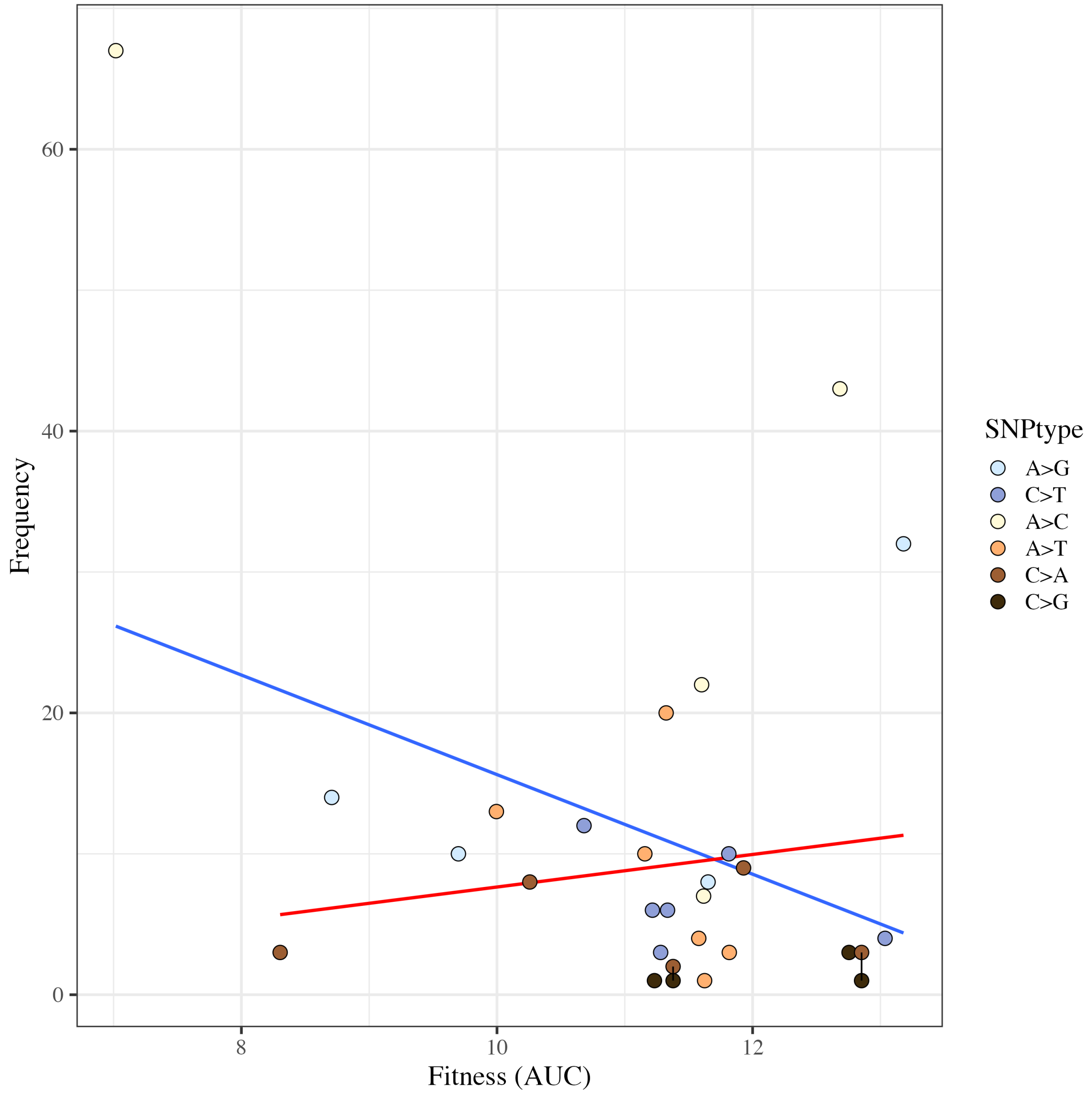

Figure S12: **Frequency of *rpoB* mutant observations across all genetic backgrounds against fitness.** Points show 29 unique mutations resulting in 27 unique AA substitutions, where 2 mutations can cause the same AA sub the points are joined with a black line. There is no significant linear relationship between fitness, as measured in minimal M9 media at 37 degrees without rifampicin, and frequency of mutant observation either including the whole data set (F = 3.74, $P$ = 0.0637 Statistical Model 9) or excluding the A>C outlier with >60 observations (AA substitution T563P) (F = 0.521, $P$ = 0.477, Statistical Model 10).

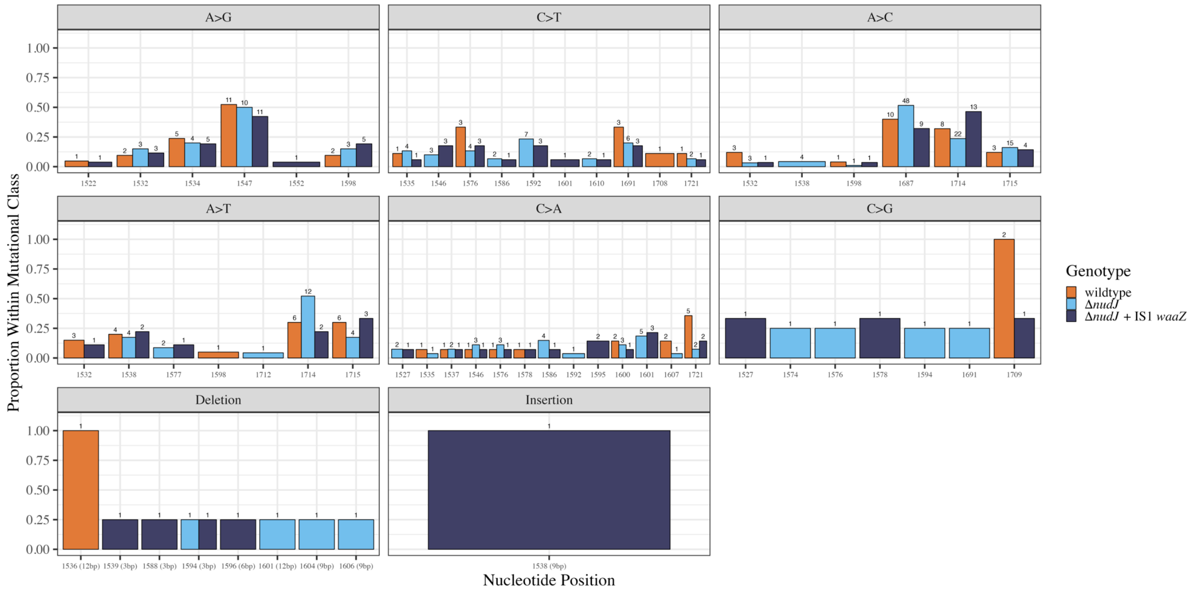

Figure S13: **Proportion of mutants accounted for by each unique mutation within each mutational class.** Bar colour indicates the genetic background in which the mutant evolved. Numbers above the bars indicate the number of sequenced mutants accounted for by the given mutation-background combination.

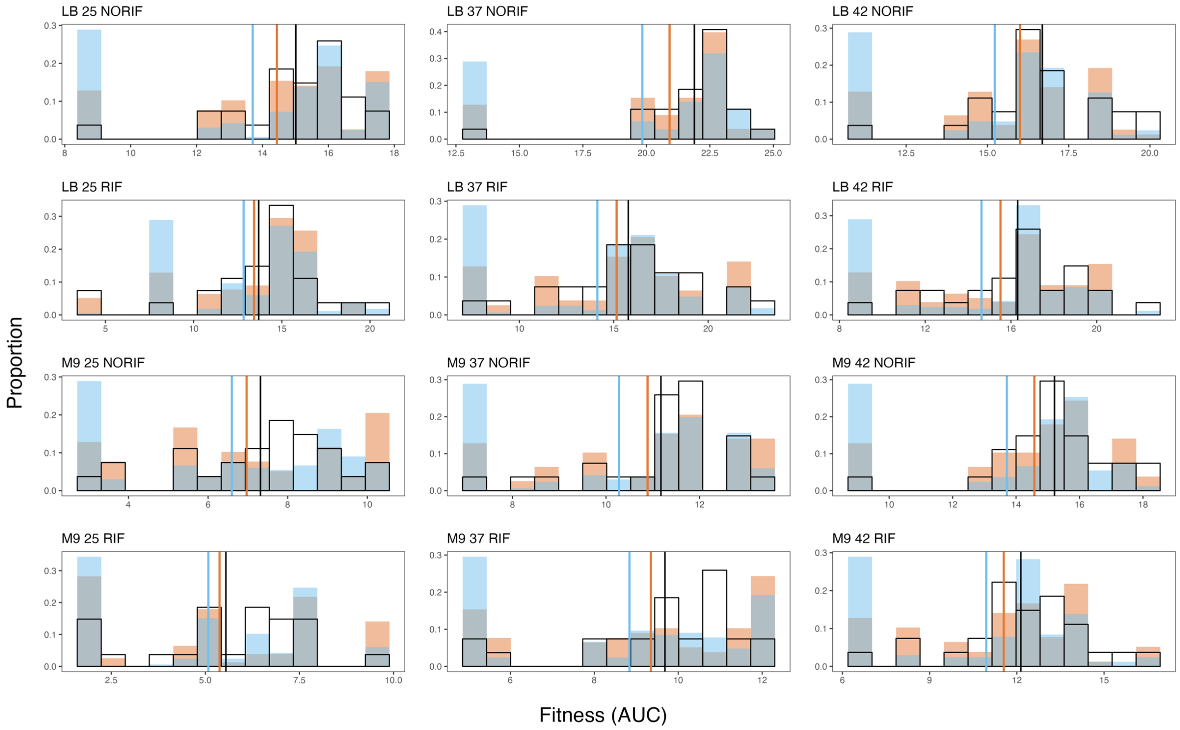

Figure S14: **DFE of rifR mutants differs by background genotype** AUC of 24 hour growth curves of rifR mutants in the MG1655 wildtype background across 12 different environments defined by media type, temperature and presence of rifampicin. The black outline shows the null DFE, with the DFE$_{\beta}$ of **Δ***nudJ* shown in blue and of the wildtype shown in orange and the means of these groups shown by vertical lines in the appropriate colour. 27 of 56 unique resistance mutations identified in this study are included.

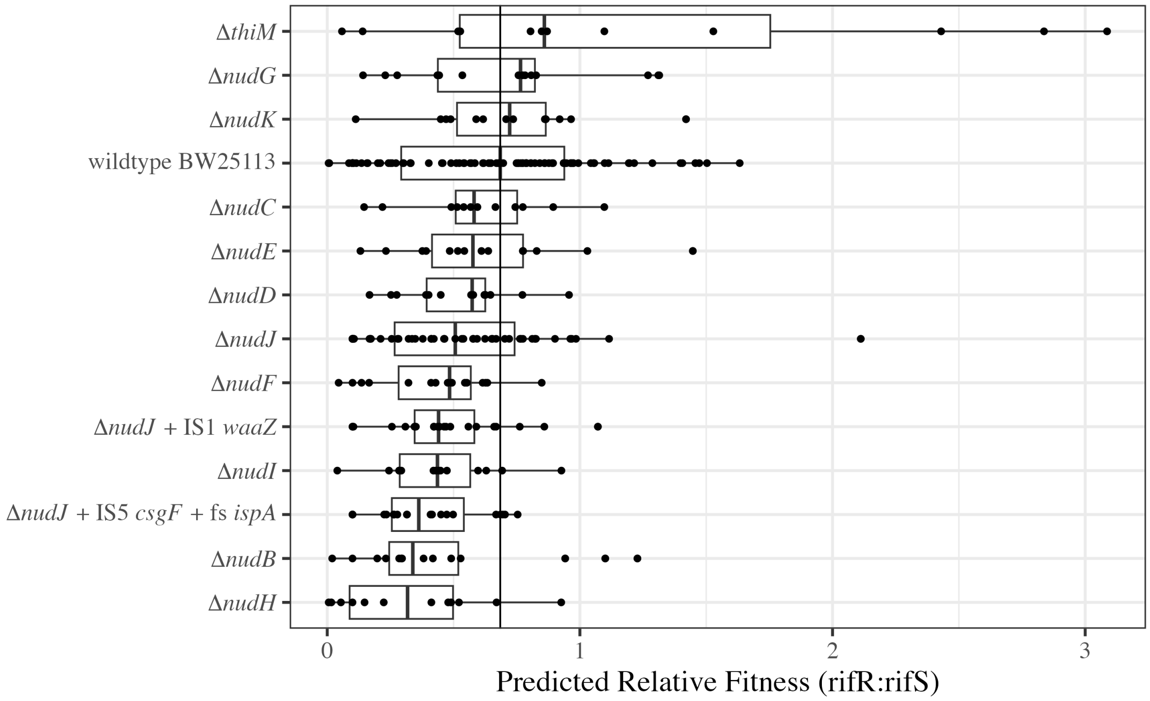

Figure S15: **Predicted cost of rifR mutants relative to rifS cells in each genetic background.** Predictions coestimated with mutational events by flan package.

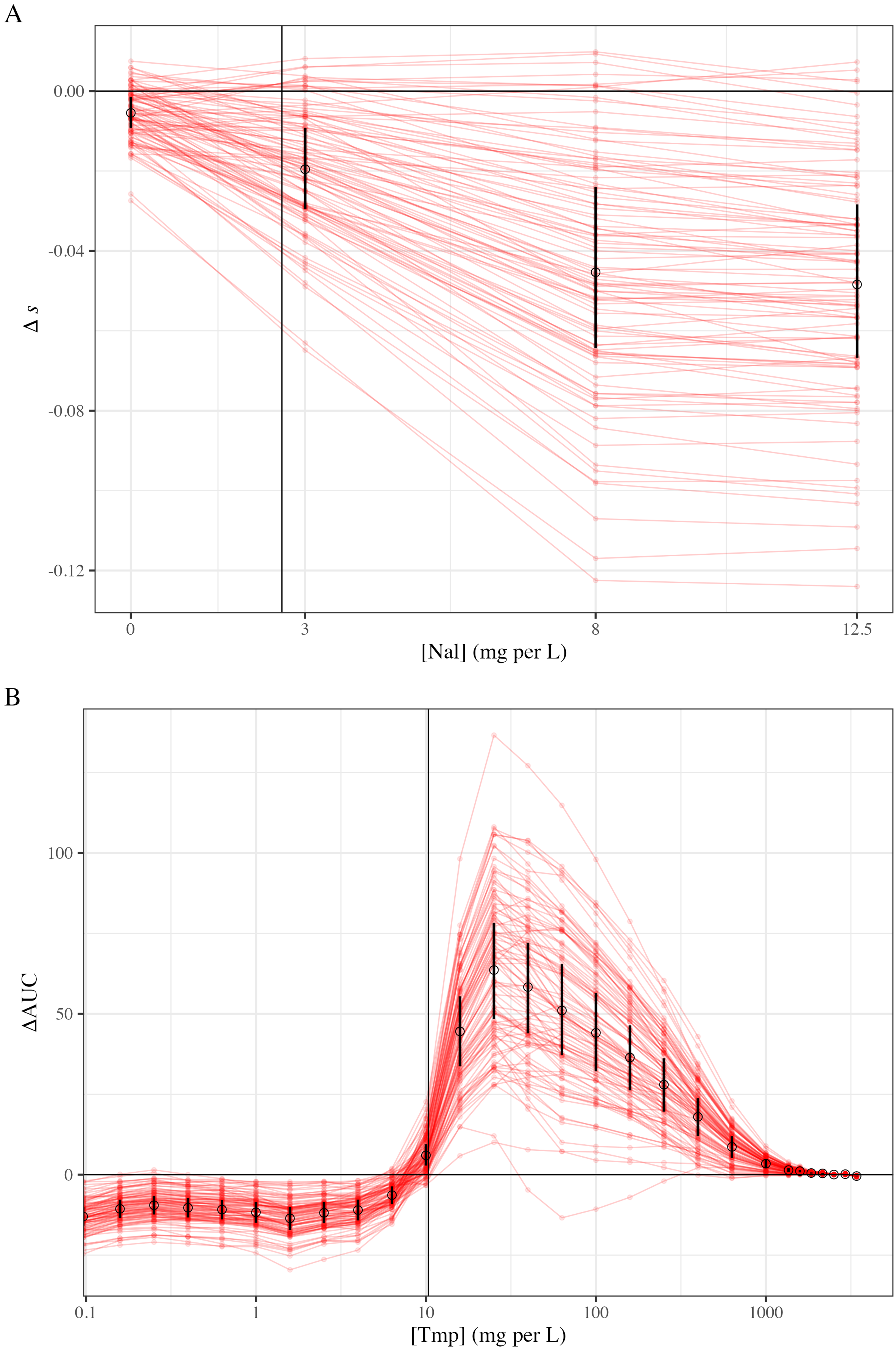

A
0.00
-0.04
-0.08
-0.12
Δ s
0
3
8
12.5
[Nal] (mg per L)
B
100
50
0
ΔAUC
0.1
1
10
100
1000
[Tmp] (mg per L)

Figure S16: **Effect of the altered mutational spectrum of Δ*nudJ* on fitness costs incurred during antibiotic adaptation.** For further details see Methods section 'Bootstrapping fitness differences …'. A) Red points connected by red lines represent the difference in mean selection coefficient (*s*) of nalidixic acid (Nal) resistant mutants accessed by **Δ***nudJ* and the wildtype (**Δ***s*) for a single bootstrap replicate. A random sample of 100 out of 1000 bootstrap replicates are shown. The Nal concentration at which *s* was measured by Harmand et al. (2017) is given on the x-axis. Black circles show the mean value of **Δ***s* across all 1000 bootstrap replicates, with the IQR across all 1000 replicates shown by black error bars. The horizontal black line shows a **Δ***s* value of of 0; points below this line indicate that mutants accessed by **Δ***nudJ* are less fit than those accessed by the wildtype and vice versa. The vertical black line represents the minimum inhibitory concentration (MIC) of Nal as given by Harmand et al. (2017) . B) As in the first panel with the following differences. The difference in average growth between mutants accessed by **Δ***nudJ* and the wildtype is given on the y-axis (**Δ**AUC). The vertical black line represents the Tmp concentration at which growth of the sensitive wildtype is suppressed to 25% of its AUC in drug-free media (IC75) as given by Palmer et al. (2015) .

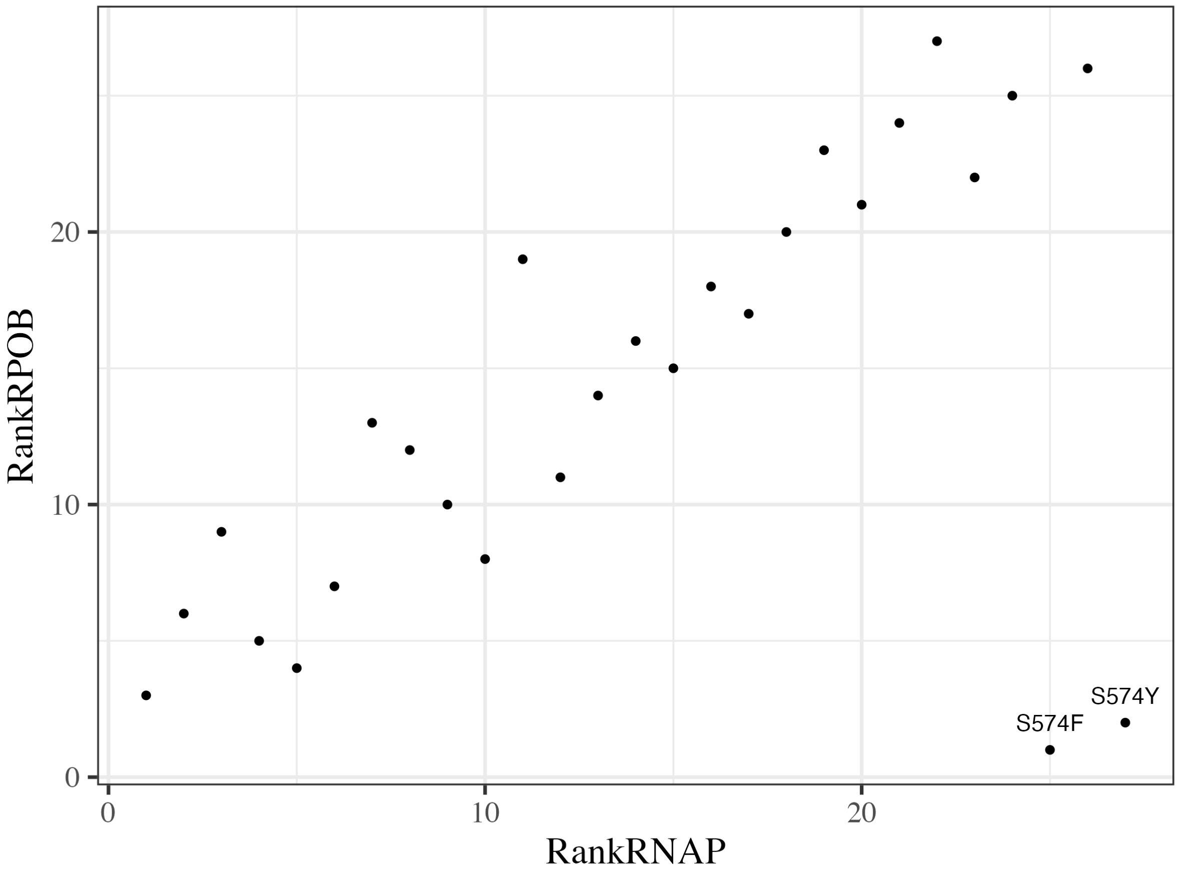

Figure S17: **Ranks of the (de)stabilising effects of *rpoB* mutants on RpoB and the RNAP complex are highly correlated.** Outliers at the 574th AA residue are labelled. Raw data can be found in Table 2. Higher ranks indicate higher values of **ΔΔ**G therefore while S574F are predicted to be highly destabilising to the RNAP complex they are the least destabilising to the RpoB subunit.

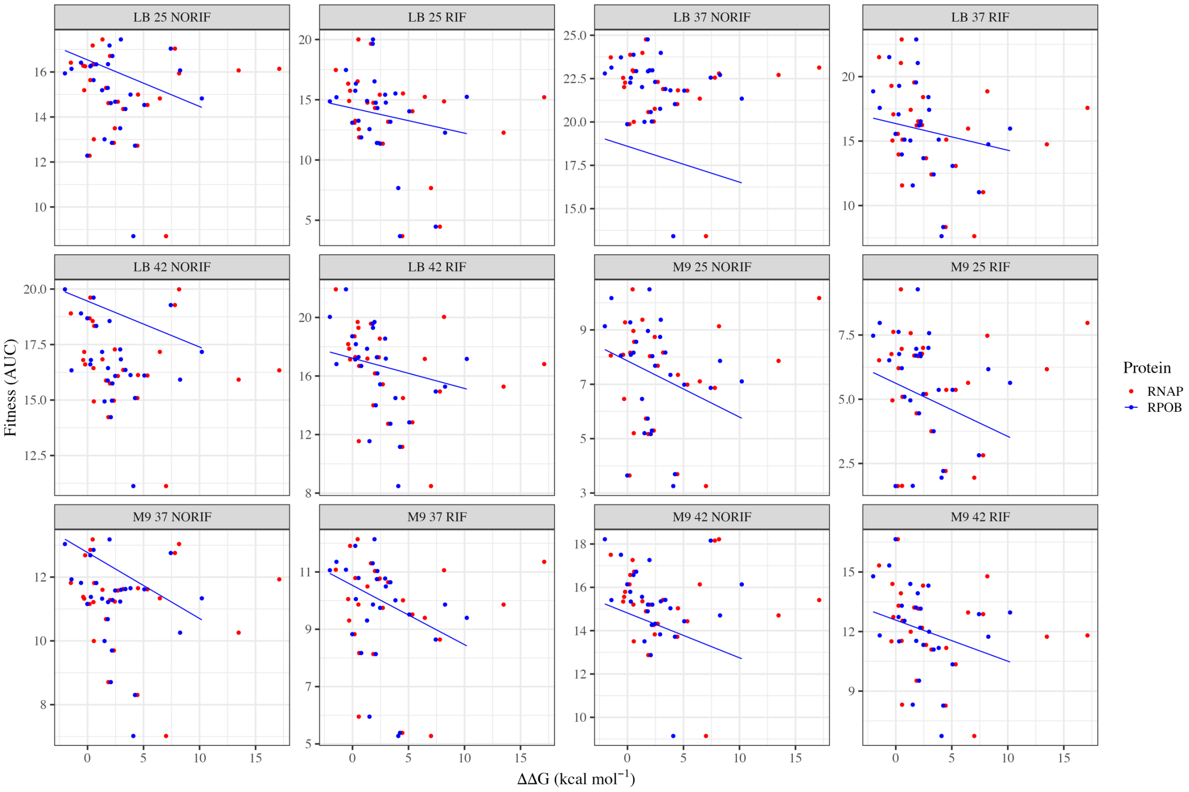

Figure S18: **Decreased stability of protein folding results in decreases in mutant fitness across 12 environments**. Fitness (measured by the proxy of AUC from 24 hour growth curves) is plotted as a function of the change in protein stability to RNAP (red) and RpoB (blue) caused by the given mutation. Greater values of **ΔΔ**G indicate reduced stability. Blue lines show predictions from Statistical Model 7. Note varying y-axis limits.

Table 1: Mutation rates across nud KOs with comparison to wildtype. All mutation rates are given as the probability of a mutational event conferring rifampicin resistance occuring in a cell division (x1e9). t statistics compare the mutation rate of the given strain to the BW25113 wildtype. Mutation rates are estimated at the mean density of all fluctuation assays.

| **Treatment** | **Rate** | **Lower** | **Upper** | **Fold Change (vs wt)** | **t-value** | **DF** | **P (Dunnett's test)** |
|---|---|---|---|---|---|---|---|
| ancestor | 21.90 | 18.40 | 26.10 | 1.000 | NA | NA | NA |
| **MG1655** | **27.70** | **24.60** | **31.20** | **1.260** | $\mathbf{3.92}$ | **283** | $\mathbf{1.74 \times 10^{-3}}$ |
| **nudB** | **12.50** | **9.41** | **16.50** | **0.568** | $\mathbf{-3.96}$ | **283** | $\mathbf{1.47 \times 10^{-3}}$ |
| nudC | 19.20 | 12.30 | 30.10 | 0.877 | $-0.573$ | 283 | 1.00 |

| | | | | | | | |
|---|---|---|---|---|---|---|---|
| nudD | 22.70 | 18.80 | 27.40 | 1.030 | 0.349 | 283 | 1.00 |
| nudE | 21.20 | 17.60 | 25.50 | 0.967 | $-0.354$ | 283 | 1.00 |
| **nudF** | **11.20** | **8.69** | **14.40** | **0.511** | $\mathbf{-5.21}$ | **283** | $\mathbf{3.65 \times 10^{-6}}$ |
| nudG | 21.50 | 17.20 | 26.90 | 0.982 | $-0.164$ | 283 | 1.00 |
| **nudH** | **4.86** | **2.28** | **10.40** | **0.222** | $\mathbf{-3.91}$ | **283** | $\mathbf{1.81 \times 10^{-3}}$ |
| nudI | 19.10 | 14.70 | 24.90 | 0.874 | $-1.01$ | 283 | 0.999 |
| **nudJ** | **6.55** | **5.42** | **7.93** | **0.299** | $\mathbf{-12.5}$ | **283** | $\mathbf{< 2.2 \times 10^{-16}}$ |
| **nudJ_ispA** | **15.90** | **12.90** | **19.50** | **0.724** | $\mathbf{-3.08}$ | **283** | $\mathbf{3.90 \times 10^{-2}}$ |
| nudJ_waaZ | 20.70 | 17.80 | 24.10 | 0.945 | $-0.727$ | 283 | 1.00 |
| nudK | 17.90 | 14.30 | 22.30 | 0.815 | $-1.83$ | 283 | 0.703 |
| thiM | 19.80 | 14.20 | 27.70 | 0.903 | $-0.599$ | 283 | 1.00 |

Table 2: All amino acid substitutions observed in the RRDR in this study. ‘AAsub’ lists the amino acid substitution in RpoB resulting from the mutation. ‘Position’ lists the mutated position in the gene, in the case of insertions and deletions the first base pair of the mutation is given. ‘SNPtype’ lists the mutational class of the mutation. ‘ancestor’, ‘nudJ’ and ‘nudJ_waaZ-IS1’ list the number of times this mutation was observed in the wildtype strain BW25113, *nudJ* deletant and the *nudJ* deletant with an IS1 insertion in *waaZ*. ‘ddG_RNAP’ and ‘ddG_RPOB’ list the predicted change in the Gibbs free energy of folding (kcal per mol) caused by the given mutation to the RNAP complex and the RpoB subunit respectively. Number of sequenced mutants is: wildtype N = 92, **Δ***nudJ* N = 201 and **Δ***nudJ*+IS1*waaZ* N = 102. ‘Ins (dup)’ refers to an insertion which is a duplication of the adjacent nucleotide sequence.

| AAsub | Position | SNPtype | ancestor | nudJ | nudJ_waaZ-IS1 | ddG_RNAP | ddG_RPOB |
|---|---|---|---|---|---|---|---|
| S508P | 1522 | A>G | 1 | 0 | 1 | 1.49 | 2.98 |
| S509R | 1527 | C>A | 0 | 2 | 1 | 0.247 | 0.537 |
| S509R | 1527 | C>G | 0 | 0 | 1 | 0.247 | 0.537 |
| L511P | 1532 | A>G | 2 | 3 | 3 | 4.51 | 3.83 |
| L511Q | 1532 | A>T | 3 | 0 | 1 | 2.71 | 2.47 |
| L511R | 1532 | A>C | 3 | 3 | 1 | 5.35 | 5.07 |

| | | | | | | | |
|---|---|---|---|---|---|---|---|
| S512P | 1534 | A>G | 5 | 4 | 5 | 1.86 | 2.08 |
| S512F | 1535 | C>T | 1 | 4 | 1 | 6.46 | 10.2 |
| S512Y | 1535 | C>A | 1 | 1 | 0 | 9.18 | 10 |
| Q513K | 1537 | C>A | 1 | 2 | 1 | -0.331 | 0.493 |
| Q513L | 1538 | A>T | 4 | 4 | 2 | 0.183 | -0.0213 |
| Q513P | 1538 | A>C | 0 | 4 | 0 | 2.06 | 4.61 |
| D516N | 1546 | C>T | 0 | 3 | 3 | 0.527 | 1.83 |
| D516Y | 1546 | C>A | 1 | 3 | 1 | -0.423 | 1.32 |
| D516G | 1547 | A>G | 11 | 10 | 11 | 0.458 | 1.96 |
| N518D | 1552 | A>G | 0 | 0 | 1 | 1.11 | 1.84 |
| T525R | 1574 | C>G | 0 | 1 | 0 | 2.43 | 2.91 |
| H526D | 1576 | C>G | 0 | 1 | 0 | 2.36 | 2.05 |
| H526N | 1576 | C>A | 1 | 3 | 1 | 1.18 | 1.44 |
| H526Y | 1576 | C>T | 3 | 4 | 3 | -1.38 | 1.2 |
| H526L | 1577 | A>T | 0 | 2 | 1 | -1.49 | -0.584 |
| H526Q | 1578 | C>A | 1 | 0 | 1 | -0.386 | 0.3 |
| H526Q | 1578 | C>G | 0 | 0 | 1 | -0.386 | 0.3 |
| R529H | 1586 | C>T | 0 | 2 | 1 | 2 | 2.22 |

| | | | | | | | |
|---|---|---|---|---|---|---|---|
| R529L | 1586 | C>A | 0 | 4 | 1 | -0.124 | -1.42 |
| S531F | 1592 | C>T | 0 | 7 | 3 | 0.592 | 0.779 |
| S531Y | 1592 | C>A | 0 | 1 | 0 | 1.34 | 6.32 |
| A532P | 1594 | C>G | 0 | 1 | 0 | 2.83 | 0.732 |
| A532E | 1595 | C>A | 0 | 0 | 2 | -0.112 | -0.729 |
| L533H | 1598 | A>T | 1 | 0 | 0 | 5 | 2.39 |
| L533P | 1598 | A>G | 2 | 3 | 5 | 2.35 | 2.18 |
| L533R | 1598 | A>C | 1 | 1 | 1 | 2.76 | 2.83 |
| G534C | 1600 | C>A | 2 | 3 | 1 | 8.42 | 6.2 |
| G534D | 1601 | C>T | 0 | 0 | 1 | 13.4 | 10.5 |
| G534V | 1601 | C>A | 0 | 5 | 3 | 13.5 | 8.26 |
| G536V | 1607 | C>A | 2 | 1 | 0 | 4.45 | 4.24 |
| G537D | 1610 | C>T | 0 | 2 | 1 | 21 | 10.2 |
| T563P | 1687 | A>C | 10 | 48 | 9 | 7.01 | 4.09 |
| P564L | 1691 | C>T | 3 | 6 | 3 | 1.65 | 1.8 |
| P564R | 1691 | C>G | 0 | 1 | 0 | 2.07 | 1.94 |
| G570S | 1708 | C>T | 1 | 0 | 0 | 11.6 | 9.47 |
| G570A | 1709 | C>G | 2 | 0 | 1 | 7.8 | 7.43 |
| L571Q | 1712 | A>T | 0 | 1 | 0 | 3.18 | 3.38 |

| | | | | | | | |
|---|---|---|---|---|---|---|---|
| I572F | 1714 | A>T | 6 | 12 | 2 | -0.3 | 1.3 |
| I572L | 1714 | A>C | 8 | 22 | 13 | -0.215 | 0.248 |
| I572N | 1715 | A>T | 6 | 4 | 3 | 0.561 | 1.52 |
| I572S | 1715 | A>C | 3 | 15 | 4 | 1.33 | 2.97 |
| S574F | 1721 | C>T | 1 | 2 | 1 | 8.17 | -2.02 |
| S574Y | 1721 | C>A | 5 | 2 | 2 | 17.1 | -1.43 |
| 12 bp del | 1536 | Deletion | 1 | 0 | 0 | | |
| 9 bp ins (dup) | 1538 | Insertion | 0 | 0 | 1 | | |
| 3 bp del | 1539 | Deletion | 0 | 0 | 1 | | |
| 3 bp del | 1588 | Deletion | 0 | 0 | 1 | | |
| 3 bp del | 1594 | Deletion | 0 | 1 | 1 | | |
| 6 bp del | 1596 | Deletion | 0 | 0 | 1 | | |
| 12 bp del | 1601 | Deletion | 0 | 1 | 0 | | |
| 9 bp del | 1604 | Deletion | 0 | 1 | 0 | | |
| 9 bp del | 1606 | Deletion | 0 | 1 | 0 | | |

Table 3: Mutational classes comparing the proportion of each SNP type in **Δ***nudJ* to the wildtype. Chi squared statistics for a binomial test of association between the genetic background and the proportion of observed mutants carrying the given mutational type are given with associated degrees of freedom and P values. P values corrected for the false discovery rate are given in the final column.

| **Mutational Class** | **Chi-Squared** | **DF** | **P** | **P (fdr corrected)** |
|---|---|---|---|---|
| A>G | 7.62000 | 2 | 0.00578 | 0.02310 |
| C>A | 0.67400 | 2 | 0.41200 | 0.65800 |
| C>G | 0.00214 | 2 | 0.96300 | 1.00000 |
| A>C | 10.70000 | 2 | 0.00107 | 0.00853 |
| A>T | 1.82000 | 2 | 0.17700 | 0.37600 |
| C>T | 0.35200 | 2 | 0.55300 | 0.73700 |
| Deletion | 1.73000 | 2 | 0.18800 | 0.37600 |
| Insertion | 0.00000 | 2 | 1.00000 | 1.00000 |

Table 4: Effects of mutational biases in **Δ***nudJ* and the wildtype on the probability of a lineage fixing via beneficial mutation. Any environment where either strain has no benefitical mutants is excluded leaving 10 environments. 'Environment' lists the media, temperature and presence or absence of rifampicin (at 50mg per L) in which fitness of the *rpoB* mutants was measured. 'AUC~b wt~' and 'AUC~b nudJ~' list the average AUC across all *rpoB* mutants observed in the wildtype or **Δ***nudJ* background respectively divided by the AUC of MG1655 in the given environment. This gives us relative growth rather than the chance of fixation which is a useful, though imperfect, proxy. '$f_{wt}$' and '$f_{nudJ}$' list the proportion of observed mutations in the given background which are beneficial (greater AUC than MG1655) in the given environment. F is the fold change in mutation rate between **Δ***nudJ* and the wildtype, this is given at 7 as observed at low densities (see figure 2). G is the ratio of '$f_{nudJ}$':'$f_{wt}$' and H is the ratio of 'AUC~b nudJ$_{\text{':'AUC}}$b wt~'. 'pFGH' defines the probability that a beneficial mutation in *rpoB* carries **Δ***nudJ* to fixation, as given in Tuffaha et al, 2023 this is calculated as the initial ratio of **Δ***nudJ* to the wildtype multiplied by F, G and H. The final column 'Ratio_AncfixesVSnJ' gives the probability of a beneficial mutation in *rpoB* carrying the wildtype to fixation relative to the

probability of a beneficial mutation in *rpoB* carrying **Δ***nudJ* to fixation in the given environment (1/pFGH)/2. The null expectation in the absence of mutational bias effects is that the final column will equal 7 (the fold change in mutation rates).

| Environment | AUCb wt | AUCb nudJ | $f$wt | $f$nudJ | F | G | H | pFGH | Ratio_AncfixesVSnJ |
|---|---|---|---|---|---|---|---|---|---|
| LB 25 NORIF | 1.05 | 1.05 | 0.397 | 0.422 | 0.156 | 1.06 | 0.996 | $8.25 \times 10^{-2}$ | 6.06 |
| LB 25 RIF | 16.4 | 15.7 | 1.00 | 1.00 | 0.156 | 1.00 | 0.956 | $7.47 \times 10^{-2}$ | 6.70 |
| LB 37 RIF | 4.14 | 3.86 | 1.00 | 1.00 | 0.156 | 1.00 | 0.933 | $7.29 \times 10^{-2}$ | 6.86 |
| LB 42 RIF | 1.72 | 1.78 | 0.872 | 0.711 | 0.156 | 0.815 | 1.03 | $6.59 \times 10^{-2}$ | 7.58 |
| M9 25 NORIF | 1.25 | 1.17 | 0.372 | 0.416 | 0.156 | 1.12 | 0.942 | $8.23 \times 10^{-2}$ | 6.08 |
| M9 25 RIF | 28.3 | 26.7 | 1.00 | 1.00 | 0.156 | 1.00 | 0.945 | $7.38 \times 10^{-2}$ | 6.77 |
| M9 37 NORIF | 1.02 | 1.02 | 0.154 | $8.43 \times 10^{-2}$ | 0.156 | 0.548 | 0.996 | $4.26 \times 10^{-2}$ | 11.7 |
| M9 37 RIF | 30.7 | 29.0 | 1.00 | 1.00 | 0.156 | 1.00 | 0.945 | $7.39 \times 10^{-2}$ | 6.77 |
| M9 42 NORIF | 1.00 | 1.00 | $1.28 \times 10^{-2}$ | $1.20 \times 10^{-2}$ | 0.156 | 0.940 | 1.00 | $7.34 \times 10^{-2}$ | 6.81 |

| | | | | | | | | | |
|---|---|---|---|---|---|---|---|---|---|
| M9 42 RIF | 1.60 | 1.65 | 0.872 | 0.711 | 0.156 | 0.815 | 1.03 | $6.58 \times 10^{-2}$ | 7.60 |
