## Supplementary Statistics for "Evolutionary potential of the *Escherichia coli* antimutator Δ*nudJ* is reduced via altered mutational spectrum"

Standard diagnostic plots are included for all linear models. All models were executed in R. Although the document can be read with no reference to the code we have chosen to also include code elements for those it might benefit. Full code as quarto documents associated with this paper are available as supplementary files.

### Model 1

The model shown in figure 1 and figure 2 of the main text fits $\log_2$ mutational events (*m*) per mL against the mean-centred $\log_2$ density (Dc), allowing for differences in intercept and slope among the 21 genotypes. Random effects of experimental plate (78 levels) nested within experimental block (21 levels) nested within experimenter (3 levels), each affecting the intercept, are also fitted. Differences in variance (i.e. heteroscedasticity) associated with estimated mutational events per mL were also included. As expected, variance was found to be negatively correlated with this variable to a power of $-0.506$.

As there are only 3 levels in the 'Experimenter' variable this may be more appropriately treated as a fixed effect, rather than a random effect. We find no improvement or significant change when modelling the experimenter effect as fixed and so the original model is retained ($AIC_{fixed}$ = 679.53, $AIC_{random}$ = 679.17, LR = 1.6, *P* = 0.2).

Predicted values from this model shown as lines of best fit in figures 2, S1, S3, S5 and S8 are calculated by dividing predicted mutational events per mL from this model by the measured value of CFU per mL; both the 'per mL' terms cancel and the result is in the units of mutational events per CFU, therefore giving a mutation rate.

```
nlme::lme(log2(m_mL)~Dc*genotype, data = FA,
             random = ~ 1 |  Experimentor/block / plate_ID,
             weights=varPower(form=~m_mL)
          )->model
```

#### ANOVA table for model 1

| | numDF | denDF | F-value | p-value |
|---|---|---|---|---|
| (Intercept) | 1 | 283 | 142.00 | <2.2e-16 |
| Dc | 1 | 283 | 358.00 | <2.2e-16 |
| genotype | 20 | 283 | 16.80 | <2.2e-16 |

| | numDF | denDF | F-value | p-value |
|---|---|---|---|---|
| Dc:genotype | 20 | 283 | 3.51 | 1.35e-06 |

### Variance and Standard Deviations of model 1

| | Variance | StdDev |
|---|---|---|
| Experimentor = | 0.0000 | 8.91e-05 |
| block = | 0.2280 | 4.78e-01 |
| plate_ID = | 0.0867 | 2.94e-01 |
| Residual | 0.3850 | 6.20e-01 |

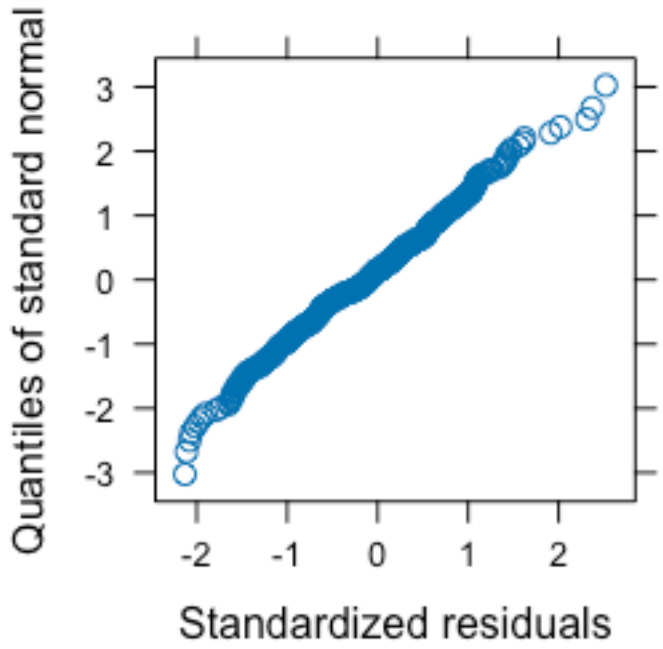


*QQ Plot of Deviance Residuals for Model 1*

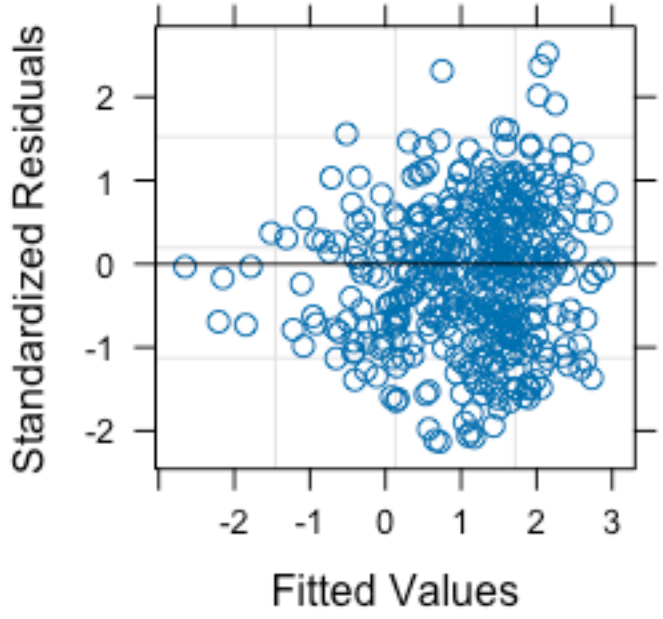


*Standardized Residuals vs Fitted Values for Model 1*

### Model 2

Model 2 fits log2 net luminesence (from an ATP based luminesence assay) against mean-centred log2 density (Dc) and genotype allowing for differences in intercept and slope among the 15 genotypes. A fixed effect of experimenter (2 levels) is also included. Random effects of experimental plate (68 levels) nested within experimental block (15 levels), each affecting the intercept, are also fitted.

```
nlme::lme(log2(LUM)~Dc*genotype+Experimentor,data=LUMdf,
          random = ~ 1 |  block / plate_ID)->model
```

#### ANOVA table for model 2

| | numDF | denDF | F-value | p-value |
|---|---|---|---|---|
| (Intercept) | 1 | 211 | 16300.00 | <2.2e-16 |
| Dc | 1 | 211 | 917.00 | <2.2e-16 |
| genotype | 14 | 211 | 13.60 | <2.2e-16 |
| Experimentor | 1 | 13 | 4.01 | 0.0667 |
| Dc:genotype | 14 | 211 | 3.03 | 0.00028 |

#### Variance and Standard Deviations of model 2

| | Variance | StdDev |
|---|---|---|

| | Variance | StdDev |
|---|---|---|
| block = | 0.3730 | 0.611 |
| plate_ID = | 0.0551 | 0.235 |
| Residual | 0.1960 | 0.442 |

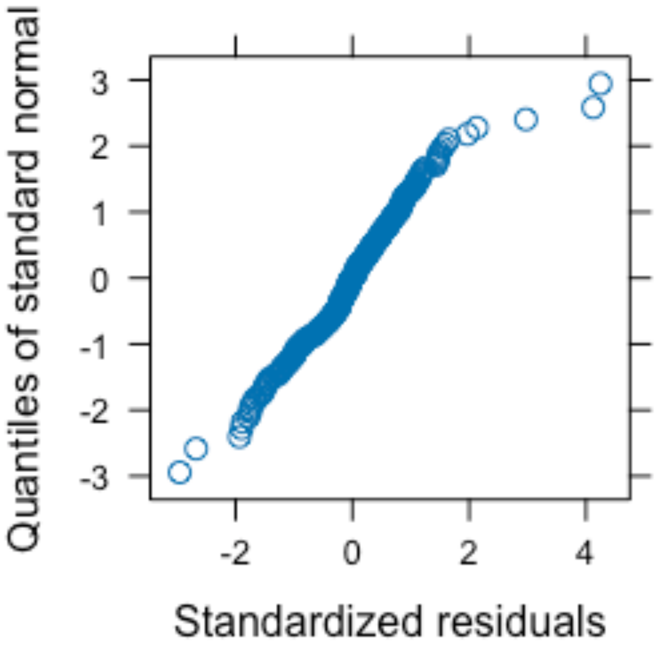


*QQ Plot of Deviance Residuals for Model 2*

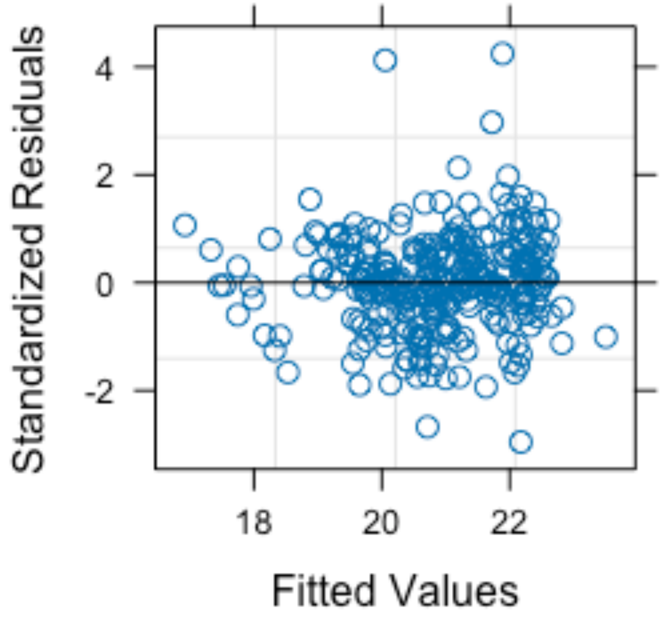


*Standardized Residuals vs Fitted Values for Model 2*

### Model 3

Model 3 fits log2 mutational events per mL against mean-centred log2 density (Dc) and genotype allowing for differences in intercept among the 3 genotypes. Differences in slope between treatments are not fitted as removing this interaction improves the model (reduced AIC). Random effects of experimental plate (6 levels) nested within experimental block (2 levels), each affecting the intercept, are also fitted. This model uses data from liquid based fluctuation assays.

```
nlme::lme(data=VBOD,log2(m_mL)~Dc+Genotype,
          random = ~ 1 |block/plate_ID)->model
```

#### ANOVA table for model 3

| | numDF | denDF | F-value | p-value |
|---|---|---|---|---|
| (Intercept) | 1 | 20 | 41.20 | 2.93e-06 |
| Dc | 1 | 20 | 48.50 | 9.32e-07 |
| Genotype | 2 | 20 | 1.18 | 0.327 |

#### Variance and Standard Deviations of model 3

| | Variance | StdDev |
|---|---|---|
| block = | 0.000 | 1.98e-05 |

| | Variance | StdDev |
|---|---|---|
| plate_ID = | 0.121 | 3.48e-01 |
| Residual | 0.335 | 5.79e-01 |

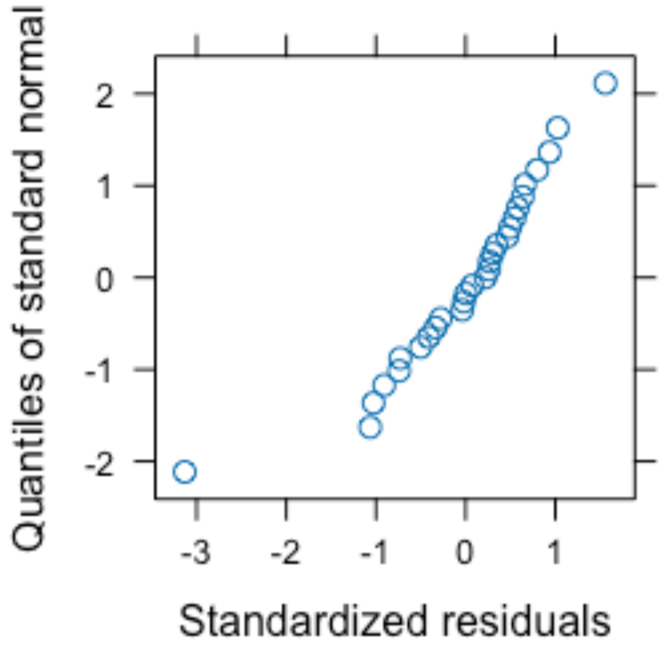


*QQ Plot of Deviance Residuals for Model 3*

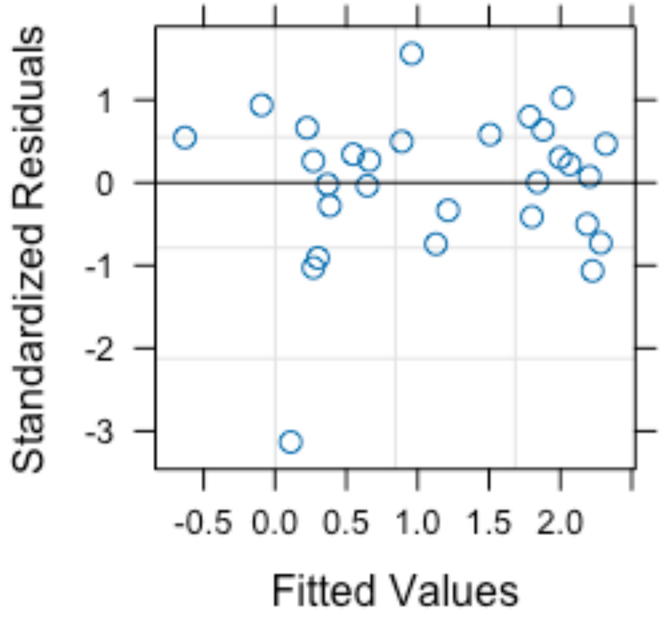


*Standardized Residuals vs Fitted Values for Model 3*

### Model 4

Model 4 fits the mutational class of rifR mutations identified in *rpoB* against the background genotype and experimental block using a multinomial regression (data shown in Figure 3). 3 genotypes and 4 experimental blocks are included with 8 possible mutational classes (transitions A>G & C>T, transversions A>C, A>T, C>A & C>G and insertions & deletions). The comparisons of mutational spectra between pairs of genotypes included in the main text are calculated by combining those two genotypes in this model and comparing this combined model back to the original. As two such comparisons are made, a false discovery rate correction is applied to the *P* values.

```
nnet::multinom(SNPtype ~ genotype+Block, data = AAtab)->model
```

#### ANOVA table for model 4

| | LR Chisq | Df | Pr(>Chisq) |
|---|---|---|---|
| genotype | 34.3 | 14 | 0.00185 |
| Block | 33.7 | 21 | 0.03860 |

*Figure 1: Probability of observing each SNP type given block and genotype as predicted by statistical model 4. 95% confidence intervals are shown.*

### Model 5

Model 5 fits the number of observed mutations in each ‘Experimental block/genotype/mutational class/unique mutation’ group as a function of mutational class, its 2-way interactions with genotype, unique mutation and block and the 3-way interaction between genotype, mutational class and unique mutation. Respectively these terms represent a test for biases in the mutational spectrum; differences in spectrum biases between genetic backgrounds; hotspots nested within mutational classes; differences in the mutational spectrum between experimental blocks; and an effect of genetic background on hotspot strengths within mutational classes. All observations are offset by the opportunity available for observing the given mutant (total number of mutants identified in the given ‘genotype/block’ group).

```
glm(formula = n ~
      genotype + SNPtype +
      SNPtype:genotype + SNPtype:UniqueMut +
      genotype:SNPtype:UniqueMut,
    family = poisson(link = "log"), data = Pois_AA)->model
```

#### ANOVA table for model 5

| | LR Chisq | Df | Pr(>Chisq) |
|---|---|---|---|
| genotype | 16.50 | 2 | $2.599 \times 10^{-4}$ |
| SNPtype | 75.70 | 7 | 1.03e-13 |
| genotype:SNPtype | 9.93 | 10 | 0.4463 |
| SNPtype:UniqueMut | 88.00 | 48 | 0.000376 |
| genotype:SNPtype:UniqueMut | 18.00 | 45 | 1 |

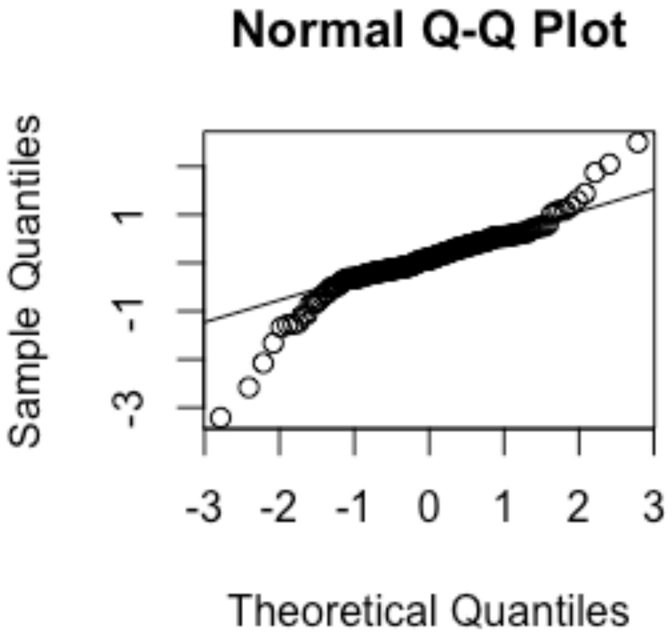


*QQ Plot of Deviance Residuals for Model 5*

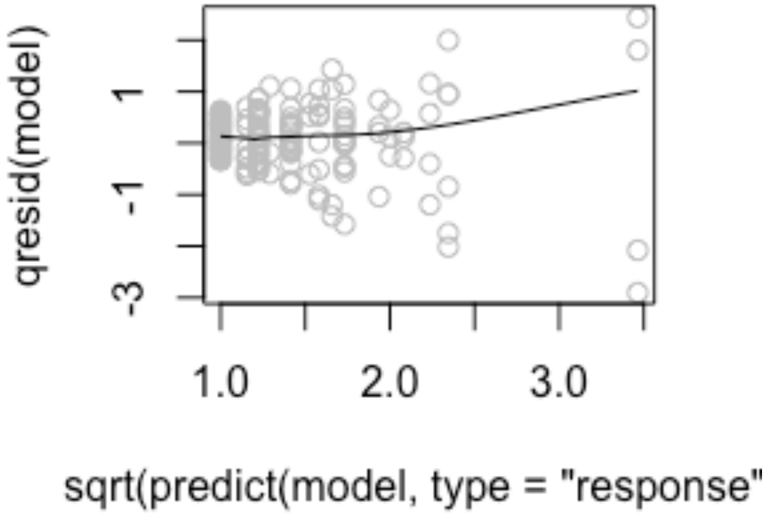


*Standardized Residuals vs Square Root of Fitted Values for Model 5*

### Model 6

Model 6, shown in figure 4 & S14, fits mean growth (measured as the area under the curve of a 24 hr growth curve) of rifR mutants in the MG1655 background against the genotype in which these mutants were observed (3 levels: **Δ***nudJ*, **Δ***nudJ* + IS1 *waaZ* & wildtype), the environment in which growth was measured, allowing for all interactions among the 3 environmental factors (antibiotic (2 levels), media (2 levels) and temperature (3 levels)), and experimental block (4 levels). Each observed 'mutant:genetic background:experimental block' set is weighted by inverse of the frequency with which it was observed. Every 'mutant:genetic background:experimental block' combination is repeated 12 times for the 12 environments in which growth was measured.

Differences in variance (i.e. heteroscedasticity) associated with both antibiotic presence and media type in which AUC was measured are also included. All possible combinations of environmental variables, either separately affecting variance or concatenated to a single variable, were tested in a model with all variable interactions, with our chosen combination selected based on AIC minimisation. The chosen model produces a significant improvement based on AIC comparison to one without heteroscedasticity (hetsced) with environmental factors ($AIC_{no\ hetsced}$ = 8563.2, $AIC_{hetsced\ temp\ +\ media}$ = 8476.5, LR = 92.7, $P < 0.0001$)

We tested fitting a model with all variable interactions including experimental block as either a fixed effect, random effect or not included. Inclusion as a fixed effect was selected based on AIC minimisation (AIC) as shown below.

| call | Model | df | AIC | BIC | logLik | Test | L.Ratio | p-value |
|---|---|---|---|---|---|---|---|---|
| Random | 1 | 38 | 8563.162 | 8766.846 | -4243.581 | | NA | NA |
| Fixed | 2 | 40 | 8556.540 | 8770.944 | -4238.270 | 1 vs 2 | 10.62137 | 0.0049385 |
| No_Block | 3 | 37 | 8569.767 | 8768.090 | -4247.883 | 2 vs 3 | 19.22630 | 0.0002455 |

```
nlme::gls(model = mean_auc ~ genotype + temperature + antibiotic +
    media + Block + temperature:antibiotic + temperature:media,
    data = DFE, weights = varComb(varFixed(~1/Freq), varIdent(form = ~1 |
        temperature), varIdent(form = ~1 | media)), method = "ML",
    control = glsControl(maxIter = 500, msMaxIter = 500))->model
```

#### ANOVA table for model 6

| | numDF | F-value | p-value |
|---|---|---|---|
| (Intercept) | 1 | 22900.00 | <2.2e-16 |
| genotype | 2 | 18.80 | 8.26e-09 |

| | numDF | F-value | p-value |
|---|---|---|---|
| temperature | 2 | 419.00 | 2.61e-146 |
| antibiotic | 1 | 175.00 | 5.07e-38 |
| media | 1 | 1410.00 | 9.07e-221 |
| Block | 3 | 5.86 | 0.000557 |
| temperature:antibiotic | 2 | 8.11 | 0.000313 |
| temperature:media | 2 | 106.00 | 8.85e-44 |

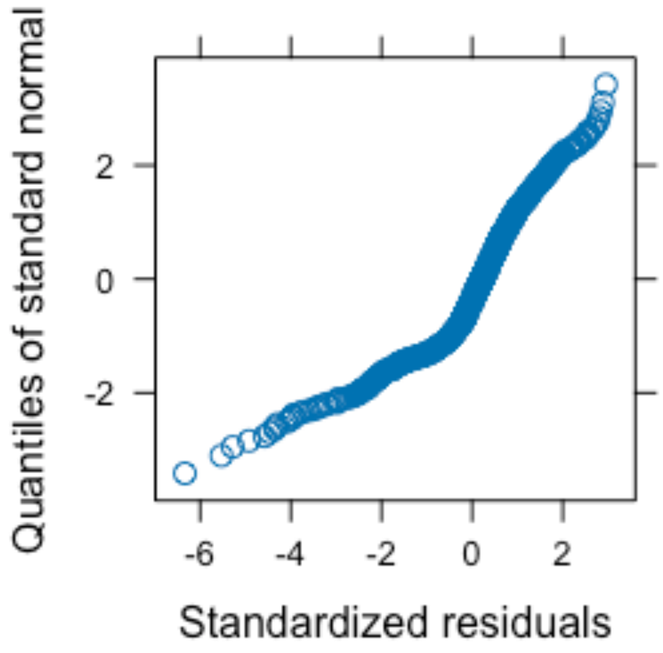


*QQ Plot of Deviance Residuals for Model 6*

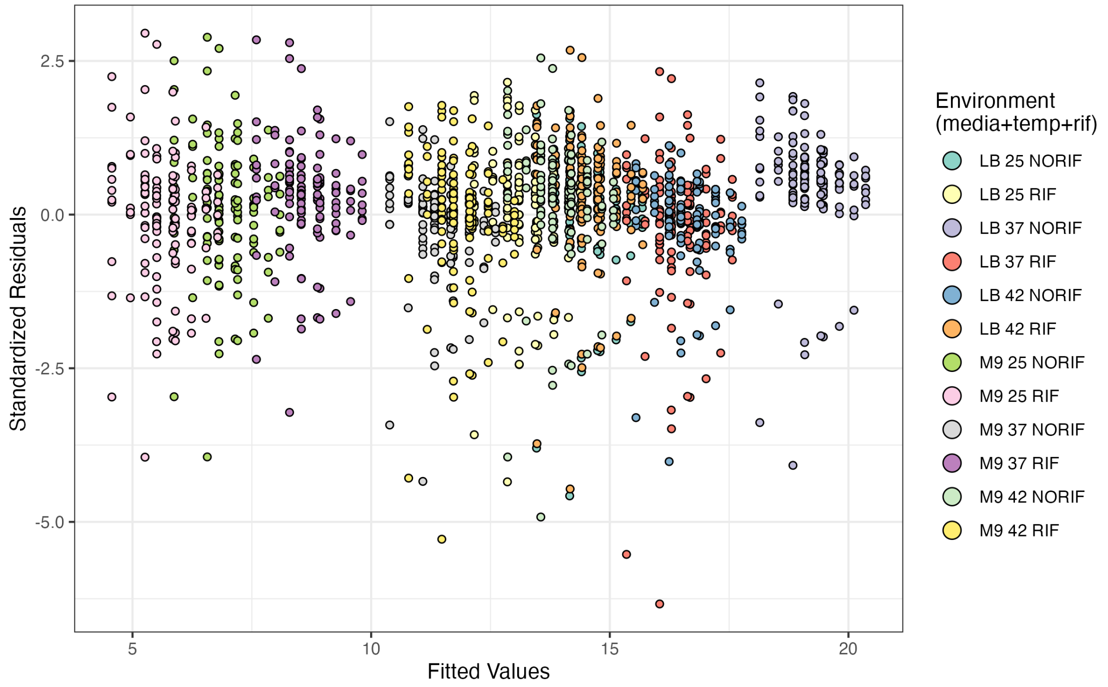


*Standardised Residuals vs Fitted Values for Model 6*

### Model 7

Model 7, shown in Fig S18, initially fits mean growth (measured as the area under the curve of a 24 hr growth curve) of rifR mutants in the MG1655 background against the **ΔΔ**G (ddG in code) to RPOB and the **ΔΔ**G (ddG in code) to the RNAP complex caused by the rifR mutation and the environment (media (2 levels), temperature (3 levels) and antibiotic (2 levels)). Each observed mutant is included once for each of the 12 environments. All interactions between explanatory variables were originally included and a minimal model, dropping the **ΔΔ**G (RNAP) term and some interactionswas, was selected based on AIC minimisation.

Differences in variance (i.e. heteroscedasticity) associated with the media type in which AUC was measured are also included. Each possible combination of environmental variables, either separately affecting variance or concatenated to a single variable and with or without an effect of the CV in growth among *rpoB* mutants in the given environment, was tested in a model with all variable interactions. 29 models were compared with the selected model chosen based on AIC minimisation. The chosen model produces a significant improvement based on AIC comparison to one without heteroscedasticity (hetsced) with environmental factors ($AIC_{\text{no hetsced}}$ = 1603.1, $AIC_{\text{hetsced media}}$ = 1562.8, LR = 42.3, $P = 7.70 \times 10^{-11}$).

```
gls(model = mean_auc ~ ddG_RPOB + temperature + media + antibiotic +
    temperature:media, data = OurMuts, weights = varIdent(form = ~1 |
    media), method = "ML", control = glsControl(maxIter = 500,
    msMaxIter = 500))->model
```

### Protein folding stability explains only a small amount of the observed variation in growth 7

*Table of summary statistics for Model 7 along with reduced models*

```
#| label: Output for ddG mod
#| echo: false
library(modelr)
AICtab2[sapply(AICtab2, is.numeric)] <- lapply(AICtab2[sapply(AICtab2,
is.numeric)], function(x) format_sci(x, digits = 3))
AICtab2%>%kable(row.names = F)
```

| Model | df | AIC | BIC | logLik | Test | L.Ratio | p-value |
|---|---|---|---|---|---|---|---|
| Null Model - retaining random effects | 3.0( | $1.96 \times 10^{3}$ | $1.97 \times 10^{3}$ | −97 | | NA | NA |
| Full Model minus fixed effect ddG_RPOB | 7.0( | $1.54 \times 10^{3}$ | $1.57 \times 10^{3}$ | −7( | 1 vs 2 | 420 | $1.47 \times 10^{-89}$ |
| Full Model including ddG_RPOB | 8.0( | $1.53 \times 10^{3}$ | $1.56 \times 10^{3}$ | −75 | 2 vs 3 | 18.7 | $1.51 \times 10^{-5}$ |

```
list(
  full = model,
  reduced = modDDG2_w_norpob
) %>%
  map_dfr(~ add_predictions(OurMuts, ., var = "yhat", type = "response"),
          .id = "model") %>%
  group_by(model) %>%
 mutate(n=nrow(OurMuts))%>%
  dplyr::summarise(
    SSE = sum((mean_auc - yhat)^2)
  )->SSEs
```

Removing the **ΔΔ**G to RpoB as an explanatory variable from this model only increases the SSE by 6.0986159% ($SSE_{full}$=2312, $SSE_{reduced}$=2453, these are the third and second models listed above respectively). This demonstrates the small effect size of this explanatory variable.

### ANOVA table for model 7

| | numDF | F-value | p-value |
|---|---|---|---|
| (Intercept) | 1 | 8160.0 | $9.44 \times 10^{-229}$ |
| ddG_RPOB | 1 | 18.9 | $1.84 \times 10^{-5}$ |
| temperature | 1 | 361.0 | $2.83 \times 10^{-54}$ |

| | numDF | F-value | p-value |
|---|---|---|---|
| media | 1 | 454.0 | $3.42 \times 10^{-63}$ |
| antibiotic | 1 | 73.2 | $5.09 \times 10^{-16}$ |
| temperature:media | 1 | 31.9 | $3.57 \times 10^{-8}$ |

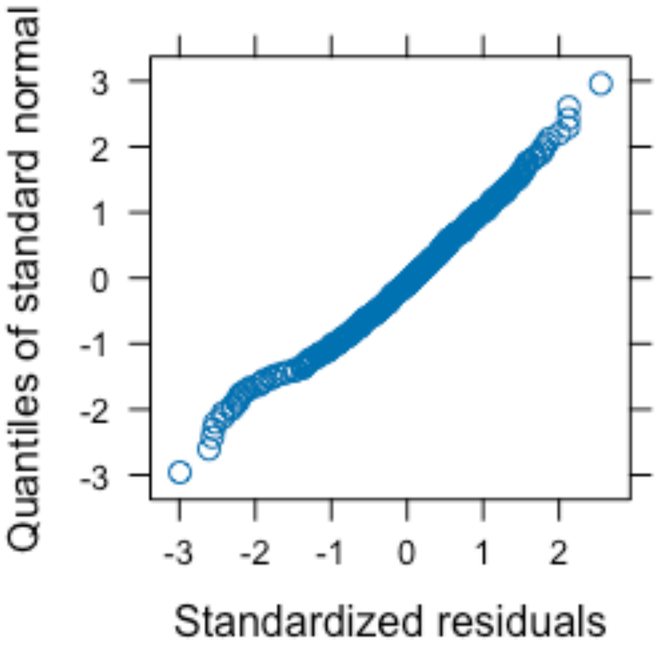


*QQ Plot of Deviance Residuals for Model 7*

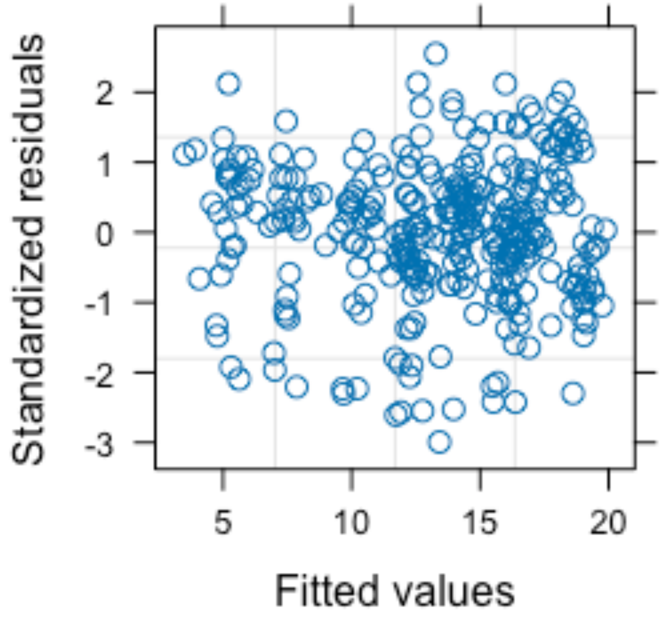


*Standardized Residuals vs Fitted Values for Model 7*

### Model 8

Estimates of carrying capacity at 125mg/L are calculated from a model fitting population density (CFU per mL) against genotype and glucose concentration centered to 125 (glcCent). This model uses only data from cultures grown in <500mg/L (within this range, population density and glucose concentration are roughly linearly related). Random effects of experimental plate (75 levels) nested within experimental block (19 levels) nested within experimenter (3 levels), each affecting the intercept, are also fitted. Differences in variance (i.e. heteroscedasticity) associated with glucose concentration were also included. Greater variance was found at higher glucose densities.

```
nlme::lme(data=FA_mono,D_genotype~glcCent+genotype,
          random=~1|Experimentor/block/plate_ID,
    weights = varPower(form = ~glucose))->model
```

#### ANOVA table for model 8

| | numDF | denDF | F-value | p-value |
|---|---|---|---|---|
| (Intercept) | 1 | 220 | 108.00 | <2.2e-16 |
| glcCent | 1 | 220 | 684.00 | <2.2e-16 |
| genotype | 14 | 220 | 6.74 | 2.21e-11 |

#### Variance and Standard Deviations of model 8

| | Variance | StdDev |
|---|---|---|
| Experimentor = | 7.31e+07 | 8550 |
| block = | 1.25e+15 | 35400000 |
| plate_ID = | 2.10e+08 | 14500 |
| Residual | 1.32e+09 | 36400 |

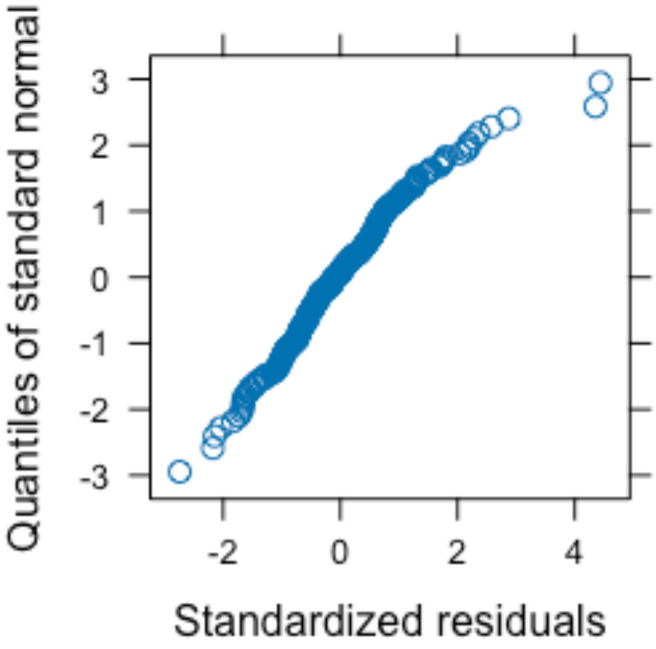


*QQ Plot of Deviance Residuals for Model 8*

Standardised Residuals vs Fitted Values for Model 8 with points coloured by glucose concentration

*Standardised Residuals vs Fitted Values for Model 8 with points coloured by glucose concentration*

### Model 9

Model 9 fits the growth of rifR mutants (measured as AUC of a 24hr growth curve of the mutant in the MG1655 background in minimal M9 media at 37 degrees in the absence of antibiotics) against frequency of observing the given mutant. An effect of SNP type interacting with the effect of frequency was initially included, the model was then selected by AIC in a stepwise algorithm.

```
lm(data=auc_oc,auc~occurences)->model
```

### ANOVA table for model 9

| | Df | Sum Sq | Mean Sq | F value | Pr(>F) |
|---|---|---|---|---|---|
| occurences | 1 | 6.88 | 6.88 | 3.74 | 0.0637 |
| Residuals | 27 | 49.70 | 1.84 | NA | NA |

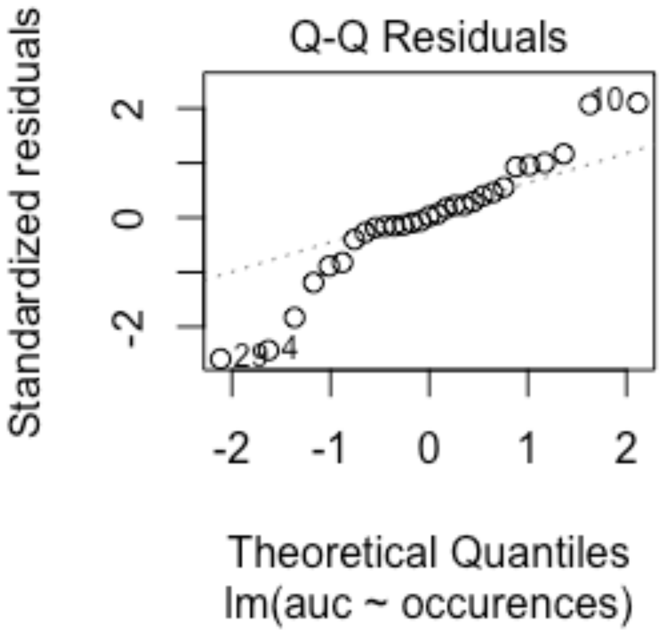


*QQ Plot of Deviance Residuals for Model 9*

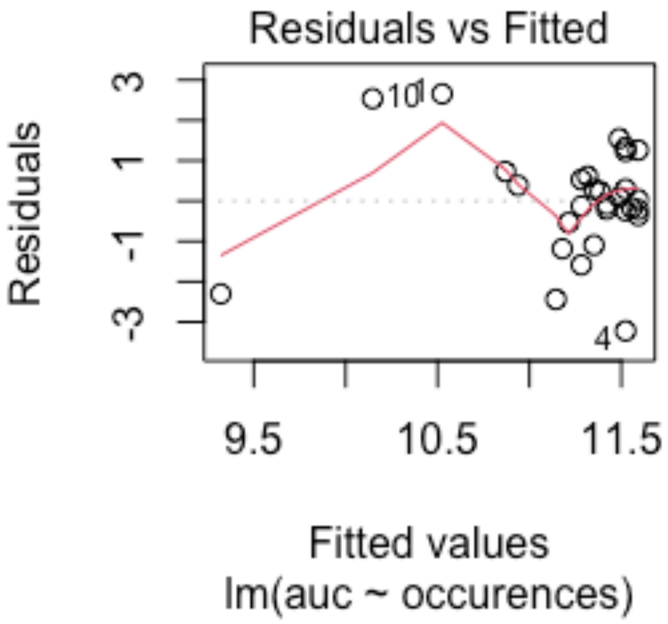


*Standardized Residuals vs Fitted Values for Model 9*

### Model 10

Model 10 fits the growth of rifR mutants (measured as AUC of a 24hr growth curve of the mutant in the MG1655 background in minimal M9 media at 37 degrees in the absence of antibiotics) against frequency of observing the given mutant. The T563P mutant, which was observed 67 times, is removed as a potential outlier. An effect of SNP type interacting with the effect of frequency was initially included, the model was then selected by AIC in a stepwise algorithm.

```
lm(data=subset(auc_oc,occurences<60),auc~occurences)->model
```

#### ANOVA table for model 10

| | Df | Sum Sq | Mean Sq | F value | Pr(>F) |
|---|---|---|---|---|---|
| occurences | 1 | 0.749 | 0.749 | 0.521 | 0.477 |
| Residuals | 26 | 37.300 | 1.440 | NA | NA |

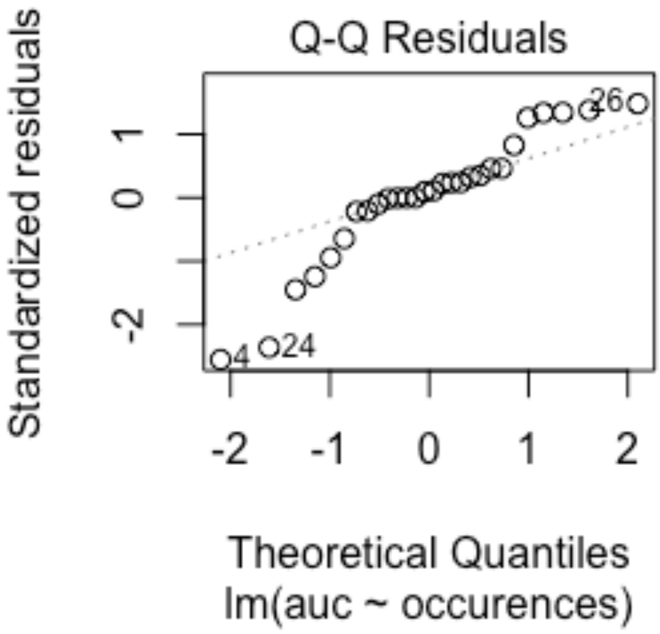


*QQ Plot of Deviance Residuals for Model 10*

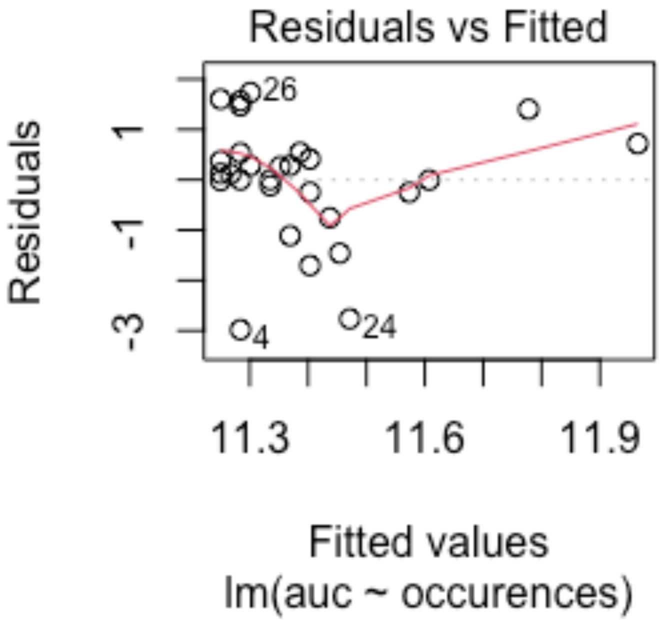


*Standardized Residuals vs Fitted Values for Model 10*
